# Evaluation and refinement of sample preparation methods for extracellular matrix proteome coverage

**DOI:** 10.1101/2020.11.25.391946

**Authors:** Maxwell C. McCabe, Lauren R. Schmitt, Ryan C. Hill, Monika Dzieciatkowska, Mark Maslanka, Willeke F. Daamen, Toin H. van Kuppevelt, Danique J. Hof, Kirk C. Hansen

## Abstract

The extracellular matrix is a key component of tissues, yet it is under-represented in proteomic datasets. Identification and evaluation of proteins in the extracellular matrix (ECM) has proved challenging due to the insolubility of many ECM proteins in traditional protein extraction buffers. Here we separate the decellularization and ECM extraction steps of several prominent methods for evaluation under real-world conditions. The results are used to optimize a two-fraction ECM extraction method. Approximately one dozen additional parameters are tested and recommendations for analysis based on overall ECM coverage or specific ECM classes are given. Compared to a standard in-solution digest, the optimized method yielded a 4-fold improvement in unique ECM peptide identifications.

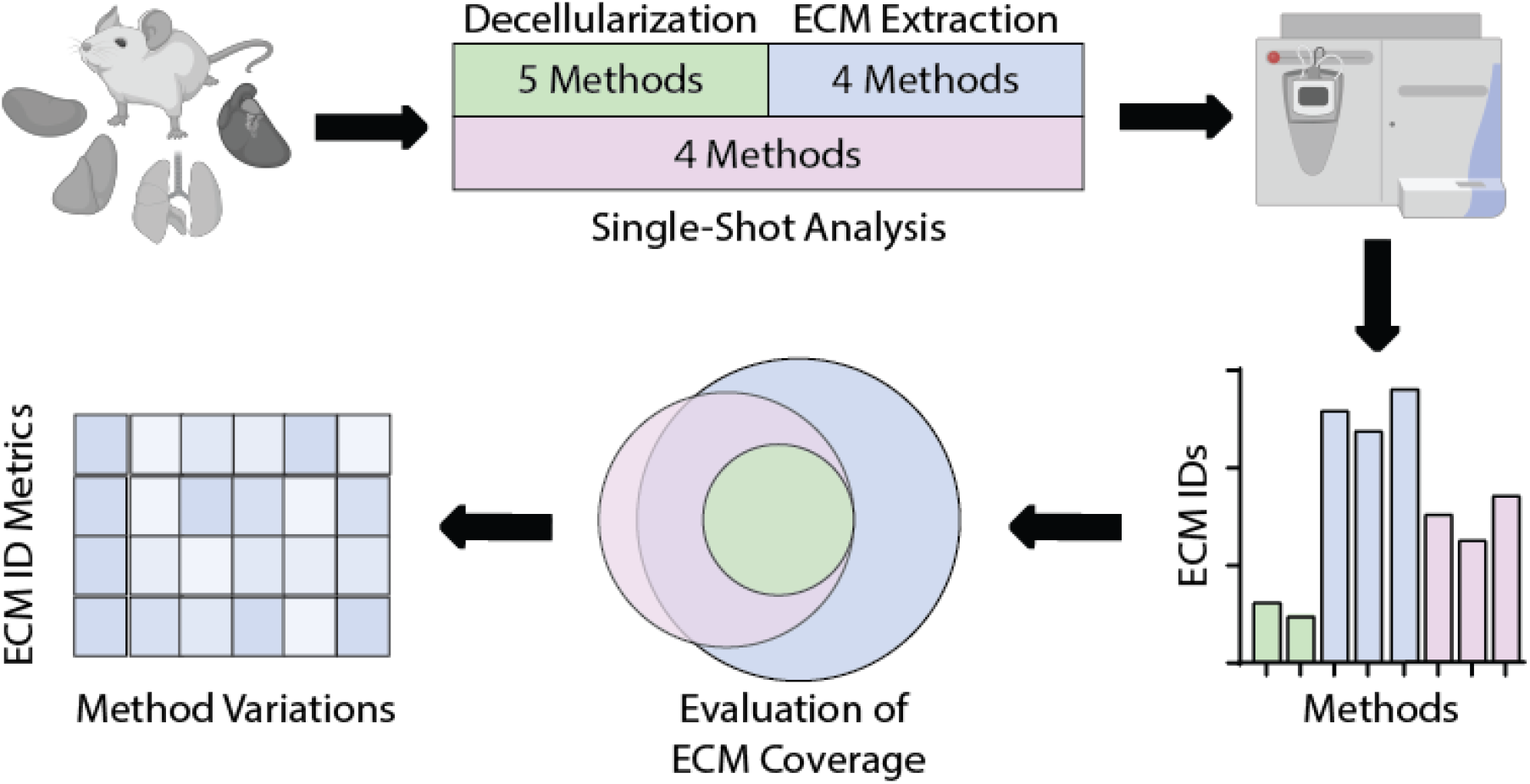

## Introduction

The extracellular matrix (ECM) is a non-cellular component of tissues which provides structural scaffolding and mediates signaling in the extracellular space to govern a wide range of biological processes including cell differentiation, wound healing, and fibrosis.^1^ Knowledge of ECM composition is critical to fields ranging from biomedical to food science, yet data regarding the ECM proteome is relatively sparse. In general, proteins of the ECM assemble into, or interact with, extended non-covalent polymers.^2^ Several core structural proteins undergo post-translational modifications including crosslink generation to further stabilize the assembled structures,^3^ rendering this sizable covalent fraction resistant to extraction in the strongest detergents and chaotropic agents.^4^ Often, collagen-containing ECM structural fibers are quantified using assays measuring hydroxyproline as a surrogate for collagen after total protein hydrolysis.^5^ The results are a crude measurement of total collagen and provide no information about collagen subtype distribution or solubility. Additionally, hydroxyproline residues are present in other cellular and extracellular proteins such as HIF-1ɑ^6^ and elastin^7^ which hinders the specificity of the assay. In other studies, second harmonic generation (SHG) and two-photon autofluorescence (TPAF) imaging have been successfully used to characterize ECM fibers by taking advantage of the intrinsic properties of collagen fibers and elastin autofluorescence to allow for analysis of ECM architecture.^8,9^ SHG and TPAF along with other imaging approaches can be used to determine the properties of ECM fibers and their degree of alignment and branching, yet they fail to provide specific qualitative and quantitative information about matrix protein abundance and subtype.

Proteomics is an attractive approach to complement these methods due to its ability to provide both compositional and quantitative readouts. Preliminary draft proteomes have been reported in recent years with deep proteome coverage of tissues obtained from protein extraction in a strong chaotrope and extended LC-MS acquisition.^10–13^ However, ECM proteins that were expected to be highly abundant (e.g. collagens I, III, V, elastin, etc.) were not found at high abundance in these datasets. This is likely a result of approximately 75-85% of the fibrillar ECM residing in a chaotrope-resistant insoluble fraction that has eluded analysis by these and other standard proteomic methods.^14^

Protein extraction protocols have been developed which specifically target enrichment of the ECM. These methods typically consist of a decellularization step followed by chaotrope extraction, either alone^15–18^ or followed by dilution and digestion with LysC and/or Trypsin in preparation for LC-MS/MS analysis.^19–25^ Biochemical methods have also been developed which use chemical digestion to solubilize and access highly insoluble ECM proteins of interest.^4,26^ We have previously developed methods which utilize chemical digestion with cyanogen bromide (CNBr)^14^ and hydroxylamine hydrochloride (HA).^4^ While both methods efficiently extract insoluble proteins, HA digestion has been the method of choice due to its safety, low cost, and lack of additional transfer steps during processing.^4^ However, non-specific Asn-X cleavages should be considered during database searching and the digestion process can induce oxidative modifications that, if extended beyond methionine single oxidation, can convolute data analysis, leaving room for improvement of the method.^4^

A wide variety of decellularization and ECM extraction methods have been published but it remains unclear how these methods perform compared to one another on a complex, ECM-rich sample. Previous matrisome enrichment method comparisons have been performed^27^ revealing the strength of chemical digestion in identifying core matrisome (structural ECM) proteins. However, significant advancements and new methods have since been developed. A direct comparison of both cell and ECM extraction methods on a whole organism sample and four additional organs serves as an important reference for future ECM proteomics. Putative proteins which compose the ECM have been previously defined using *in silico* and proteomic approaches, generating the MatrisomeDB which is divided into core matrisome and matrisome-associated proteins.^28^ Here, we utilize core matrisome annotations of collagens, ECM glycoproteins, and proteoglycans for comparison of ECM protein characterization. For this comparison, five widely used decellularization methods^4,19,29^ and four methods for single-shot extraction and analysis^21,30,31^ (Table 1) have been used for evaluation. In addition, four ECM extraction methods: a two-step extraction with guanidine hydrochloride followed by hydroxylamine hydrochloride (Gnd-HCl/HA) digestion,^4^ chaotrope-assisted in-solution digest with ultrasonication (CAISU),^19^ chaotrope-assisted in-solution digest (CAIS),^20^ and surfactant and chaotropic agent assisted sequential extraction/on-pellet digestion (SCAD)^21^ were evaluated. The findings are used to make recommendations for tissue analysis based on factors including matrisome protein sequence coverage, the number of matrisome proteins identified, and variability of results.

**Table 1.** Descriptions of tested decellularization, ECM extraction, and single shot methods with required time and composition of each extraction buffer.

| Method | Steps Involving Buffer Exchange | Time | Buffer Composition |
| --- | --- | --- | --- |
| 1-Fraction A<sup>4</sup> | 3 | 3 hrs | 1. 50mM Tris-HCl, 0.25% CHAPS, 25mM EDTA 3M NaCl, 10μL/mL PI, pH 7.4 |
| 1-Fraction B<sup>28</sup> | 5 | 7 days | 1. 10mM Tris-HCl, 0.1% EDTA, 1x PI, pH 8.0<br>2. 50mM Tris-HCl pH 7.4, 10mM MgCl<sub>2</sub>, 1% benzonase, 1x PI |
| 2-Fraction | 2 | 2 hrs | 1. 150mM NaCl, 50mM Tris-HCl, 5% glycerol, 1% NP-40, 1mM MgCl<sub>2</sub>, 1x PI<br>2. 50mM Tris-HCl pH 7.4, 2% CHAPS, 2mM EDTA, 3M NaCl, 1x PI |
| 3-Fraction<sup>18</sup> | 4 | 3 hrs | 1. 150mM NaCl, 50mM Tris-HCl pH 7.5, 5% glycerol, 1% NP-40, 1mM MgCl<sub>2</sub>, 1x PI, 1% benzonase<br>2. 150mM NaCl, 50mM Tris-HCl pH 7.5, 5% glycerol, 1% NP-40, 0.5% DOC, 0.1% SDS, 1x PI, 1% benzonase<br>3. 500mM NaCl, 50mM Tris-HCl pH 7.5, 5% glycerol, 1% NP-40, 2% DOC, 1% SDS, 1x PI, 1% benzonase |
| 4-Fraction | 10 | 8.5 hrs | 1. Buffer C (HEPES pH7.9, MgCl<sub>2</sub>, KCl, EDTA, sucrose, glycerol, NaOV), 1x PI<br>2. Buffer W (HEPES pH7.9, MgCl<sub>2</sub>, KCl, EDTA, sucrose, glycerol, NaOV), 1x PI<br>3. Buffer N (HEPES pH7.9, MgCl<sub>2</sub>, KCl, EDTA, glycerol, DOC, NP-40, NaOV), 1x PI, 1% benzonase<br>5. Buffer M (HEPES pH7.9, MgCl<sub>2</sub>, KCl, EDTA, sucrose, glycerol, DOC, NP-40, NaOV), 1x PI<br>6. Buffer CS (Pipes pH6.8, MgCl<sub>2</sub>, NaCl, EDTA, sucrose, SDS, NaOV), 1x PI |
| Gnd-HCl/HA<sup>4</sup> | 3 | 2.5 days | 1. 6M Gnd-HCl, 100mM ABC pH8.0<br>2. 1M HA, 4.5M Gnd-HCl, 0.2M K<sub>2</sub>CO<sub>3</sub>, pH 9.0 |
| CAISU<sup>17</sup> | 1 | 19 hrs | 1. 6M Gnd-HCl<br>2. 10mM Tris-HCl pH8.5, 2M Gnd-HCl, 2.7M Urea, 3% ACN<br>3. 10mM Tris-HCl pH8.5, 600mM Gnd-HCl, 800mM Urea, 3% ACN |
| CAIS<sup>18</sup> | 1 | 1 day | 1. 8M Urea, 100 mM ABC pH8.0<br>2. 2M Urea, 100mM ABC pH8.0, 1000U PNGaseF |
| SCAD<sup>20</sup> | 3 | 1.7 days | 1. 4% SDS, 50 mM Tris<br>2. 8M Urea, 100mM ABC, pH 8.0 |
| In-Gel<sup>30</sup> | 11 | 21 hrs | 1. 25mM ABC, 50% ACN<br>4. 25mM/50mM ABC |
| In-Solution | 1 | 17 hrs | 1. 25mM ABC |
| SPEED<sup>29</sup> | 1 | 17 hrs | 1. TFA<br>2. 2M Tris Base |

## Methods

### Tissue Preparation

Whole Mouse Powder (WMP) Production: Whole male C67BL/6J mouse was frozen in liquid nitrogen and fractured into approximately 1 cm^3^ pieces. Fur and blood were not removed from the mouse before milling. The resulting pieces were kept frozen in liquid nitrogen before milling to a fine powder under liquid nitrogen using a SPEX 6870 freezer/mill. The milled powder was kept frozen and lyophilized for 24 hours. Approximately 100 mg aliquots of lyophilized powder were then delipidated by 4 successive extractions with 2 mL 100% ice-cold acetone and briefly dried at room temperature in a fume hood. Isolated mouse organs (heart, liver, kidney, and lung) were prepared using the same method.

### Decellularization Methods

#### 1-Fraction A^4^

Approximately 5mg of WMP was homogenized (Bullet Blender, Model BBX24, Next Advance, Inc.) for 3 minutes on power 8 in 200 μL/mg of high salt buffer (50 mM Tris-HCl, 0.25%CHAPS, 25 mM EDTA, 3 M NaCl, pH 7.4) supplemented with 10 μL/mL fresh protease inhibitor (Halt™ Protease Inhibitor Cocktail, Thermo Scientific™ #78429) with the addition of approximately 50 1mm glass beads. Homogenate was vortexed at 4°C for 20 minutes. Homogenized tissue was spun at 18,000 x g (4 °C) for 15 min. The resulting supernatant was removed, and the pellet was further extracted with 1 ml high salt buffer two times with homogenization after each buffer addition. Cellular extracts were pooled into a single soluble fraction.

#### 1-Fraction B^29^

Approximately 5 mg of WMP was placed in 200 μL/mg 10 mM Tris-HCl (pH 8.0) with 0.1% ethylenediamine tetraacetic acid (EDTA) and 1X protease inhibitors (Halt™ Protease Inhibitor Cocktail, Thermo Scientific™ #78429) at 4°C for 48 h. Triton X-100 (Sigma-Aldrich #T9284) was added to a final concentration of 3% and samples were vortexed at medium power at 4°C for 72 h. The solution was changed every 24 h by spinning at 18,000 x g (4 °C) for 15 min and samples were resuspended by vortexing. The resulting supernatant from each buffer exchange was pooled into a single cellular fraction. Samples were then incubated in 50 mM Tris-HCl pH 7.4, 10 mM MgCl_2_, 1% benzonase (Millipore #70746) at 37°C for 24 h, spun at 18,000 x g (4 °C) for 15 min, and the supernatant was discarded. Decellularized WMP was then washed with PBS with 1X protease inhibitors for 24 h to remove residual reagents. All steps were conducted under continuous shaking.

#### 2-Fraction

Approximately 5 mg of lyophilized WMP was homogenized (Bullet Blender, Model BBX24, Next Advance, Inc.) in 200 μl/mg buffer 1 (150 mM NaCl, 50 mM Tris–HCl (pH 7.5), 5% glycerol, 1% NP-40 (US Biological #N3500), 1 mM MgCl_2_, 1X protease inhibitors (Halt™ Protease Inhibitor Cocktail, Thermo Scientific™ #78429)) at power 8 for 3 minutes with the addition of approximately 50 1mm glass beads and vortexed at 4°C for 20 minutes. Homogenized tissue was spun at 18,000 x g (4 °C) for 15 min and the supernatant was collected (fraction 1). After the addition of each extraction buffer, samples were resuspended for 1 minute at power 8 using the Bullet Blender. Sample was then homogenized in 200 μL/mg (of starting tissue dry weight) buffer 2 (50mM Tris-HCl (pH 7.4), 2% CHAPS, 2mM EDTA, 3M NaCl, 1X protease inhibitors) at power 8 for 3 minutes and vortexed at 4°C for 15 minutes. Homogenized tissue was spun at 18,000 x g (4 °C) for 15 min and the supernatant was collected (fraction 2).

#### 3-Fraction^19^

Approximately 5 mg of lyophilized WMP was homogenized (Bullet Blender, Model BBX24, Next Advance, Inc.) in 500 μl PBS containing protease inhibitors (Halt™ Protease Inhibitor Cocktail, Thermo Scientific™ #78429) at power 8 for 3 minutes with the addition of approximately 50 1mm glass beads. Homogenate was centrifuged at 16,000 x g for 20 min at 4 °C and the supernatant was collected. After the addition of each extraction buffer, samples were resuspended for 1 minute at power 8 using the Bullet Blender. Sample was then homogenized in 500 μL buffer 1 (150 mM NaCl, 50 mM Tris–HCl (pH 7.5), 5% glycerol, 1% NP-40 (US Biological #N3500), 1 mM MgCl_2_, 1X protease inhibitors, 1% benzonase (Millipore #70746)) at power 8 for 15 seconds before incubating 20 mins on ice. Homogenate was centrifuged at 16,000 x g for 20 min at 4 °C and the supernatant was collected and pooled with the first wash to make fraction 1. Sample was then homogenized in 500 μL buffer 2 (150 mM NaCl, 50 mM Tris–HCl (pH 7.5), 5% glycerol, 1% NP-40, 0.5% sodium deoxycholate (DOC), 0.1% SDS, 1X protease inhibitors, 1% benzonase) at power 8 for 15 seconds before incubating 20 mins on ice. Homogenate was centrifuged at 16,000 x g for 20 min at 4 °C and the supernatant was collected (fraction 2). Sample was then homogenized in 500 μL buffer 3 (500 mM NaCl, 50 mM Tris–HCl (pH 7.5), 5% glycerol, 1% NP-40, 2% DOC, 1% SDS, 1X protease inhibitors, 1% benzonase) at power 8 for 15 seconds before incubating 20 mins at room temperature. Homogenate was centrifuged at 16,000 x g for 20 min at 4 °C and the supernatant was collected (fraction 3). Soluble fractions were precipitated with 80% acetone and resuspended in 8M urea for subsequent digestion.

#### 4-Fraction (Millipore Compartment Protein Extraction Kit, #2145)

5 mg of lyophilized WMP was homogenized (Bullet Blender, Model BBX24, Next Advance, Inc.) in 500 μl of Buffer C containing protease inhibitors (Halt™ Protease Inhibitor Cocktail, Thermo Scientific™ #78429) for 3 minutes on power 8 with the addition of approximately 50 1mm glass beads. Homogenate was vortexed at power 4 for 20 min at 4 °C. Homogenate was then centrifuged at 16,000 x g for 20 min at 4 °C and the supernatant was collected (fraction 1) and flash frozen. After the addition of each extraction buffer, samples were resuspended for 1 minute at power 8 using the Bullet Blender. The pellet was resuspended in 400 μl of Buffer W containing protease inhibitors and vortexed at power 4 for 20 min at 4°C. Sample was centrifuged at 16,000 x g for 20 min at 4 °C and the supernatant was discarded. The pellet was then resuspended in 150 μl of Buffer N containing protease inhibitors, 1% benzonase (Millipore #70746) and vortexed at power 4 for 20 min at 4°C. Homogenate was centrifuged at 16,000 x g for 20 min at 4 °C and the supernatant was collected (fraction 2) and flash frozen. Centrifugation was repeated and the remaining supernatant was added to the N fraction. The pellet was resuspended in 400 μl of Buffer W containing protease inhibitors and vortexed at power 4 for 20 min at 4°C. Sample was centrifuged at 16,000 x g for 20 min at 4 °C and the supernatant was discarded. The pellet was then resuspended in 100 μl of Buffer M containing protease inhibitors and vortexed at power 4 for 20 min at 4°C. Homogenate was centrifuged at 16,000 x g for 20 min at 4 °C and the supernatant was collected (fraction 3) and flash frozen. The pellet was then resuspended in 200 μl of Buffer CS containing protease inhibitors and vortexed at power 4 for 20 min at 4 °C. 9. Homogenate was centrifuged at 16,000 x g for 20 min at 4 °C and the supernatant was collected (fraction 4). The pellet was resuspended in 150 μl of Buffer C containing protease inhibitors and vortexed at power 4 for 20 min at 4 °C. Homogenate was then centrifuged at 16,000 x g for 20 min at 4 °C and the supernatant was pooled with fraction CS before flash-freezing the CS fraction. Additional washes were performed by resuspending the pellet in 500 μl of PBS containing protease inhibitors and vortexing at power 4 for 5 min at 4°C. Homogenate was centrifuged at 16,000 x g for 20 min at 4°C and the supernatant was discarded. Washes were repeated three times.

#### Digestion and Preparation of Extracts from Decellularization for MS

All decellularization fractions were digested using the filter aided sample preparation (FASP) protocol as previously described^32^ using 10 kDa molecular weight cutoff filters (Sartorius, Vivacon #VN01H02). Digests were performed for 16 hours in 25 mM ABC pH 8.0 using trypsin at a 1:100 enzyme:protein ratio at 37°C in an oven. Digested samples were desalted using Pierce™ C18 Spin Tips (Thermo Scientific #84850) according to the manufacturer’s protocol.

### Single-Shot Methods

#### Surfactant and Chaotropic Agent Assisted Sequential Extraction/On-pellet Digestion (SCAD)^21^

Approximately 5 mg of WMP was solubilized in 300 μL of buffer (4% SDS, 50 mM Tris buffer) and incubated at 95 °C for 10 min. After allowing the solution to return to room temperature, protein extract was reduced with 10 mM dithiothreitol (DTT) for 30 min at room temperature and alkylated with 50 mM iodoacetamide (IAM) for an additional 15 min in the dark. The reaction was then quenched with an additional 2mM DTT. SDS was removed by two rounds of precipitation. For the first precipitation, cold acetone (−20 °C) was added to a final concentration of 80% (v/v), and the protein was precipitated overnight at −20 °C. For the second round, 80% acetone/water (v/v) was added, followed by incubation at −20 °C for 2 h. The samples were centrifuged at 18,000 × g for 15 min, and the pellet was briefly air-dried in a fume hood. Pellet was dissolved in 125uL of 8M urea and on-pellet digestion was performed with Lys-C (1:100, Wako #121-05063) for 4 h at 37 °C. Samples were diluted with 875uL of 50mM Tris buffer along with Trypsin (1:100, Promega #V511C) for overnight digestion. The reaction was quenched with 1% FA. Digested samples were desalted using Pierce™ C18 Spin Tips (Thermo Scientific #84850) according to the manufacturer’s protocol.

#### In-Gel Digest^31^

Approximately 1 mg of WMP was suspended in SDS-PAGE loading buffer (5% SDS, 250 mM Tris-HCl pH 7.0, 50% glycerol) and heated to 95°C for 5 minutes. The protein homogenate was then loaded onto a 3-8% TAE gel and run 1 cm into the gel. The gel was stained with Brilliant Blue R (Sigma-Aldrich #B7920) and the entire protein-containing band was excised. In-gel digestion was performed as previously described.^31^ Digested samples were desalted using Pierce™ C18 Spin Tips (Thermo Scientific #84850) according to the manufacturer’s protocol.

#### In-Solution Digest

Approximately 1 mg of WMP was suspended in 20uL of 50mM ABC, 0.2% ProteaseMax. Sample was vortexed for 60 minutes. Sample was diluted with 100uL 50mM ABC. DTT was added to the sample to reach a final concentration of 10mM and incubated at 37°C for 15 minutes. Sample was removed from heat and allowed to cool for 5 minutes to condense and then spun down to collect condensate. IAM was added to sample at 2.5 molar excess DTT and incubated in the dark at room temperature for 30 minutes. Alkylation was quenched with 10% excess DTT. Digestion was performed using 3ug Trypsin in 0.03% ProteaseMax overnight at 37°C. Samples were then acidified to 0.1% FA to stop digestion. Digested samples were desalted using Pierce™ C18 Spin Tips (Thermo Scientific #84850) according to the manufacturer’s protocol.

#### Sample Preparation by Easy Extraction and Digestion (SPEED)^30^

Approximately 5 mg of WMP was suspended in 100 μL trifluoroacetic acid (TFA) and incubated at room temperature for 10 min. Samples were further irradiated for 10 s at 800 W using a microwave oven. Samples were neutralized with 2M TrisBase using 10x volume of TFA before adding DTT to 10 mM and reducing for 30 minutes at 37°C. Iodoacetamide was added to a 2.5x molar excess over DTT and samples were incubated in the dark for 15 minutes. Digestion was carried out for 20 hrs at 37°C using trypsin at an enzyme:protein ratio of 1:50. Digested samples were desalted using Pierce™ C18 Spin Tips (Thermo Scientific #84850) according to the manufacturer’s protocol.

### ECM Extraction Methods

#### Hydroxylamine Chemical Digest (Gnd-HCl/HA)^4^

ECM-enriched pellets were homogenized in 6M guanidine hydrochloride (Gnd-HCl), 100mM ammonium bicarbonate (ABC) at power 8 for 1 minute (Bullet Blender, Model BBX24, Next Advance, Inc.) and vortexed (power 5) at room temperature overnight. Homogenate was spun at 18,000 x g (4 °C) for 15 min and the supernatant was collected as the Gnd-HCl fraction. Remaining pellets were reduced and alkylated by incubating in 10 mM DTT, 100 mM ABC pH 8.0 for 30 minutes at 37°C before adding 2.5x molar excess of IAM (over DTT) and incubating in the dark for 15 minutes. Samples were spun at 18,000 x g (4 °C) for 15 min and the supernatant was discarded. Pellets were then treated with freshly prepared hydroxylamine (HA) buffer (1 M NH_2_OH−HCl, 4.5 M Gnd−HCl, 0.2 M K_2_CO_3_, pH adjusted to 9.0 with NaOH) at 200 μL/mg of the starting tissue dry weight. Each tube was placed under a stream of nitrogen gas and sealed before being homogenized at power 8 for 1 minute and incubated at 45 °C with shaking (1000 rpm) for 4 h. Following incubation, the samples were spun for 15 min at 18 000 x g, and the supernatant was removed and stored as the hydroxylamine (HA) fraction at −80°C until further proteolytic digestion with trypsin. All Gnd-HCl and HA fractions were subsequently subjected to enzymatic digestion with trypsin using a filter aided sample preparation (FASP) approach^32^ and desalted using Pierce™ C18 Spin Tips (Thermo Scientific #84850) according to the manufacturer’s protocol.

#### Chaotrope-assisted In-solution Digest with Ultrasonication (CAISU)^19^

Digestion of ECM-enriched pellets was performed in two steps. Samples were resuspended in 6M guanidinium hydrochloride (Gnd-HCl). Samples were reduced with 10mM DTT for 30 minutes at room temperature and alkylated with 50mM IAM for 15 minutes in the dark and the reaction was quenched with additional DTT. The first digestion was done at 37°C for 2 hrs with LysC (1:50 enzyme to protein ratio) in 10 mM Tris–HCl (pH 8.5) containing 2M Gnd-HCl, 2.7M urea, and 3% acetonitrile. Prior to the two-hour incubation, samples were sonicated for 15 minutes (37°C) using a Bioruptor plus ultrasonicator (Diagenode). The second digestion step was done using fresh LysC (1:50 enzyme to protein ratio) and trypsin (1:20 enzyme to protein ratio) in 600 mM Gnd-HCl, 800 mM urea, and 3% acetonitrile at 37°C overnight. An additional sonication step was also performed prior to the overnight digest. Digested samples were desalted using Pierce™ C18 Spin Tips (Thermo Scientific #84850) according to the manufacturer’s protocol.

#### Chaotrope-assisted In-solution Digest (CAIS)^20^

ECM-enriched pellets were resuspended in 50 μL of 8M urea, 100 mM ABC, 10 mM DTT and incubated with continuous agitation at 1,400 rpm for 2 hrs at 37 °C. Samples were cooled to RT and 500 mM IAM in water was added to a final concentration of 25 mM. Samples were then incubated in the dark for 30 min at RT. Sample buffer was diluted to 2M urea with 100 mM ABC pH 8.0 and 1000U PNGaseF was added before incubating with continuous agitation at 1,400 rpm for 2 hr at 37 °C. Lys-C (1 μg) was added and samples were incubated with continuous agitation at 1,400 rpm for 2 hr at 37 °C. Trypsin (3 μg) was added and samples were incubated with continuous agitation at 1,400 rpm O/N at 37 °C. A second aliquot of trypsin (1.5 μg) was added and samples were incubated with continuous agitation at 1,400 rpm for an additional 2 hrs at 37 °C. Trypsin was then inactivated by acidifying the sample with 10% formic acid. Acidified samples were centrifuged at 16,000 x g for 5 min at RT and the supernatant was collected. Digested samples were desalted using Pierce™ C18 Spin Tips (Thermo Scientific #84850) according to the manufacturer’s protocol.

#### SCAD^21^

Protocol was performed as described above using WMP pellets decellularized using the 1-fraction A method rather than WMP.

#### Sample Preparation of Isolated Organs for Instrument Comparison

Samples of isolated heart, kidney, and liver from C57BL/6J mice were dissected and flash frozen by the Jackson Lab. Each organ was cryo-milled into a fine, homogenous powder using a mortar and pestle under liquid nitrogen and lyophilized for future processing. Organ samples from 3 individual mice were pooled and mixed thoroughly before weighing. Samples were decellularized using the 1-fraction A method, as described above, followed by the Gnd-HCl/HA, CAISU, CAIS, and SCAD ECM extraction methods. All ECM extraction methods were performed as previously described for WMP comparisons.

### ECM Extraction Method Optimization

#### Gnd-HCl/HA Digest Optimization

For antioxidant testing, hydroxylamine digest was performed as described above with the addition of 50 mM methionine, 100 μM/500 μM caffeic acid, 100 μM/500 μM gallic acid, or 2 mM ascorbic acid (vitamin C) to the hydroxylamine digest buffer immediately before pellet treatment. Acid pretreatment testing was performed by incubating the insoluble pellet in 0.2% formic acid (FA) for 10 minutes before spinning at 18,000 x g (4 °C) for 15 min and discarding the supernatant. For each condition, HA buffer at the stated hydroxylamine concentration was pH-adjusted prior to pellet treatment and the digest was carried out for the stated time under vortexing.

#### PNGase F Digestion

PNGase F digestion was performed on a 10 kDa cutoff filter (Sartorius #VN01H02) prior to FASP digestion. Filters were first equilibrated with successive washes of 0.1% FA followed by 8M urea, 100 mM ABC pH 8.0. All washes were spun through at 14,000 x g for 15 minutes. Samples were then loaded onto filters and spun at 14,000 x g for 20 minutes before washing with 200 μL of 8M urea, 100 mM ABC. Filters were then washed with 3 aliquots of 100 μL 50 mM ABC pH 8.0 before adding 1000U PNGase F (NEB #P0704) in 2M urea, 100 mM ABC pH 8.0 directly to the top of the filter membrane. PNGase F digestion was allowed to proceed for 2 hours at 37°C before spinning at 14,000 x g for 15 minutes to remove digest volume, retaining deglycosylated protein on the membrane. Filters were then washed with 100 μL 50 mM ABC pH 8.0 to remove residual buffer. Trypsin digest was performed in 20 mM ABC pH 8.0 with 0.02% ProteaseMax (Promega #V207A) for 16 hours using a 1:100 enzyme:protein ratio. Samples were eluted and acidified in 150 μL 0.2% FA. Digested samples were desalted using Pierce™ C18 Spin Tips (Thermo Scientific #84850) according to the manufacturer’s protocol.

#### PNGase F Digestion with GAG Removal

Removal of N-linked glycans and glycosaminoglycans (GAGs) using PNGaseF (NEB #P0704), heparinase II (NEB #P0736), and chondroitinase ABC (Sigma-Aldrich #C3667) was performed across multiple digestion steps. Filter equilibration, sample loading, and pre-digest washes were performed as described above for the PNGase F digestion. After performing three washes with 50 mM ABC, 1000U PNGase F and 0.5U heparinase II in 100 μL 100 mM NaCl, 20 mM Tris-HCl, 1.5 mM CaCl_2_ was added directly to each filter membrane. Digests were carried out for 16 hours at 37°C before spinning for 15 minutes at 14,000 x g to remove digest volume, retaining deglycosylated protein on the membrane. The filter was then washed with 100 μL 50 mM Tris-HCl pH 7.5 before adding 100 μL of 50 mM Tris-HCl pH 7.5, 60 mM sodium acetate, 0.02% BSA containing 0.5U chondroitinase ABC. Chondroitinase ABC digestion was performed for 8 hours at 37°C before spinning filters for 15 minutes at 14,000 x g to remove digest volume. Filters were then washed with 100 μL 50 mM ABC pH 8.0. Trypsin digest was performed by adding trypsin at a 1:100 enzyme:protein ratio in 100 μL 25 mM ABC pH 8.0, 0.02% ProteaseMax (Promega #V2071) and allowing the digest to proceed for 16 hours at 37°C. Samples were eluted and acidified in 150 μL 0.2% FA. Digested samples were desalted using Pierce™ C18 Spin Tips (Thermo Scientific #84850) according to the manufacturer’s protocol.

### Data Acquisition and Processing

#### Sample Quantification and Normalization

Samples from all methods were quantified after trypsin digest using a Micro BCA protein assay kit (Thermo Fisher Scientific #23235) according to the manufacturer’s protocol. Aliquots containing 10 μg of protein from each digest were loaded onto PierceTM C18 Spin Tips (Thermo Scientific #84850) and desalted using the manufacturer’s protocol. Each sample was dried to approximately 2 μL under vacuum and was brought up in 16 μL 0.1% FA.

#### MS/MS Acquisition

Global proteomics for all comparative method testing was carried out (n=3 per group) on an LTQ Orbitrap Velos mass spectrometer (Thermo Fisher Scientific) coupled to an Eksigent nanoLC-2D system through a nanoelectrospray LC − MS interface. Eight μL of each sample was injected into a 20 μL loop using the autosampler. The analytical column was then switched on-line at 600 nl/min over an in house-made 100 μm i.d. × 150 mm fused silica capillary packed with 2.7 μm CORTECS C18 resin (Waters; Milford, MA). After 10 min of sample loading at 600 nL/min, each sample was separated on a 120-min gradient consisting of a linear gradient from 2-8% ACN with 0.1% formic acid (FA) at a flow rate of 600 nL/min from 3 minutes to 20 minutes, followed by a linear gradient from 8-22% ACN with 0.1% FA at a flow rate of 350 nL/min from 20 minutes to 90 minutes. Gradient elution was followed by a linear increase to 60% ACN at 350 nL/min from 90 minutes to 98 minutes and further to 90% ACN from 98 minutes to 104 minutes to remove remaining peptides. The column was then re-equilibrated with 2% ACN in 0.1% FA at 350 nL/min from minutes 104 to 120. LC mobile phase solvents consisted of 0.1% formic acid in water (Buffer A) and 0.1% formic acid in acetonitrile (Buffer B, Optima™ LC/MS, Fisher Scientific, Pittsburgh, PA). Data acquisition was performed using the instrument supplied Xcalibur™ (version 4.1) software. The mass spectrometer was operated in the positive ion mode. Each survey scan of m/z 300–2,000 was followed by collision-induced dissociation (CID) MS/MS of the 20 most intense precursor ions with an isolation width of 2.5 m/z. Dynamic exclusion was performed after fragmenting a precursor 2 times within 15 sec for a duration of 30 sec. Singly charged ions were excluded from CID selection. Normalized collision energies of 35 eV were employed using helium as the collision gas.

timsTOF Pro data acquisition for isolated organs was carried out (n=3 per group) using a Bruker NanoElute LC system through a nanoelectrospray LC−MS interface. One μL of each sample was injected into a 20 μL loop using the autosampler. The analytical column was then switched on-line at 600 nl/min over a 15cm NanoElute column (Bruker Daltonics) using ReproSil 1.9 μm C18 resin (Dr. Maisch GmbH, Germany). After 4 min of sample loading at 800.0 bar, each sample was separated on a 120-min gradient. For cellular and Gnd-HCl fractions, the LC method consisting of a linear gradient from 2-24% ACN with 0.1% formic acid (FA) at a flow rate of 500 nL/min from 2 minutes to 112 minutes, followed by a linear increase to 95% ACN at 500 nL/min from 112 minutes to 115 minutes. Column washing at 95% ACN was performed from minutes 115 to 120. For HA fractions, the same LC method was followed but with a linear gradient from 2-20% ACN from 2 minutes to 112 minutes. LC mobile phase solvents consisted of 0.1% formic acid in water (Buffer A) and 0.1% formic acid in acetonitrile (Buffer B, Optima™ LC/MS, Fisher Scientific, Pittsburgh, PA). Data acquisition was performed using the manufacturer-supplied otofControl (version 6.0) software with the instrument default data-dependent acquisition (DDA) parallel accumulation–serial fragmentation (PASEF) method with a cycle time of 1.1s. The mass spectrometer was operated in the positive ion mode. In brief, each survey scan of m/z 100–1,700 was followed by 10 PASEF MS/MS scans employing collision-induced dissociation (CID). Active exclusion was performed with an intensity threshold of 2500 cts/s and releasing after 0.2 min, reconsidering precursors if the current intensity is 4-fold greater than the previous intensity.

#### Data Processing

All Orbitrap-acquired raw MS files were converted to .mgf format using Proteome Discoverer (Thermo Fisher Scientific) using the default parameters. Converted files were then searched using an in-house Mascot™ server (Version 2.5, Matrix Science). For Orbitrap acquired data mass tolerances were ± 15ppm for MS peaks, and ± 0.6 Da for MS/MS fragment ions. Protein probability thresholds were set at 99.9% with a minimum of 2 peptides and peptide thresholds were set at 95% using local false discovery rate (LFDR) scoring implemented in Scaffold (version 4.9.0, Proteome Software Inc.), resulting in a protein FDR of 0.0% and a peptide FDR of 0.5%. For timsTOF Pro acquired raw files, searching was performed using Peaks Studio (Version 10.5, Bioinformatics Solutions Inc.). Data refinement was performed by correcting precursor mass only, associating features with chimera scans, and filtering features for charge between 2 and 8. PeaksDB search was performed using ± 15ppm for MS peaks, and ± 0.1 Da for MS/MS fragment ions. Data was filtered to 1% FDR at the peptide level and protein probability threshold was set to P ≤ 0.01 using PEAKS.

For all searches, data was searched against SwissProt (17,029 sequences) restricted to *Mus musculus* using version 1.1 of the CRAPome for common contaminants.^33^ Trypsin specific cleavage was used in searches for cellular and enzyme-extracted ECM fractions, while HA/Trypsin specificity was used for hydroxylamine digested fractions, both allowing for 2 missed cleavages. HA/Trypsin specificity was defined as cleaving C-terminal of K and R but not before P, as well as C-terminal of N but not before C, F, H, I, M, N, Q, S, W, or Y based on previous HA cleavage data.^4^ Fixed modifications were set as carbamidomethyl (C). Variable modifications were set as oxidation (M), oxidation (P) (hydroxyproline), Gln->pyro-Glu (N-term), deamidated (NQ), and acetyl (Protein N-term).

To identify additional experimentally-induced modifications, Mascot searches were performed as described above with additional variable modifications (Search 1: Oxidation (D), Oxidation (HW), Oxidation (K), Oxidation (R); Search 2: Carbamyl (K), Carbamyl (R), Carbamyl (N-term)).

#### Data Analysis

##### Protein Classification

Core matrisome annotations, including collagens, proteoglycans, and ECM glycoproteins, were mapped using the mouse matrisome database (MatrisomeDB, http://matrisomeproject.mit.edu/).^28^ Cellular proteins are defined as all proteins not annotated within the MarisomeDB as core matrisome or matrisome-associated.

##### Normalization to Equal Run Time

To compare methods generating different numbers of fractions, PSMs were normalized to total run time by dividing the sum of PSMs in all fractions of a single method by the number of fractions analyzed.

##### Weighted Average Sequence Coverage

Sequence coverage was determined using an in-house Mascot™ server (Version 2.5, Matrix Science) and Scaffold (version 4.9.0, Proteome Software Inc.). For each round of sample runs, fractional abundance of a given protein was calculated by dividing peptide spectral matches for that protein by the total number of peptide spectral matches for proteins in that category (i.e. collagens) across all compared samples. The sequence coverage for an individual protein was then multiplied by the fractional abundance. The weighted sequence coverage for proteins in that category were summed to provide the weighted average sequence coverage for that category.

##### Statistical Significance

P-values were calculated using a two-tailed equal variance student’s t-test. P-values in text are reported as the least significant relevant comparison. In figures, “*” denotes p<0.05, “**” denotes p<0.01, and “***” denotes p<0.001.

##### Overall Method Scoring

Overall method scores were determined by scoring each method based on a variety of relevant criteria and calculating a weighted average of these scores. Each category is scored from 1 to 10, with 1 representing the lowest possible score and 10 representing the highest. For categories “Time” and “ECM in Cellular Fraction”, higher scores are given to methods with lower actual values. Decellularization was scored based on time (10%), ease (10%), MS compatibility of buffers (10%), total protein identification (30%), precision (20%), and ECM in the cellular fraction (20%). For comparison to decellularization methods, single-shot methods were compared on all metrics except ECM in the cell fraction, dividing the weight for this metric evenly among the other categories. ECM extraction methods were scored based on time (10%), ease (10%), collagen identification (30%), glycoprotein and proteoglycan identification (30%), and precision (20%). Identical scoring and weighting were applied to single-shot methods for comparison to ECM extraction methods.

## Results

### Evaluation of Cellular Protein Extraction Methods for ECM Enrichment

To compare the various decellularization methods, we evaluated extraction of cellular proteins and preservation of ECM proteins for subsequent ECM analysis. Whole mouse powder (WMP) was chosen for method development as it is a complex mixture that includes a representative sampling of all tissues and covers matrix components across a wide dynamic range. The single-shot methods were also compared for extraction of cellular proteins (Figure 1A). For decellularization methods with multiple fractions, each fraction was analyzed separately, and data is presented as both fraction sums and per-fraction averages. When normalized to equal MS run time, the 1-fraction A method generates more cellular peptide spectral matches (PSMs) than other methods, with 15% more PSMs than the next-highest 4-fraction method (p=0.014), while in-solution digest yields the least (Figure 1B). No significant differences in cellular spectral matches were observed between 2-fraction, 3-fraction, and 4-fraction extractions when comparing per-fraction averages. Of note, the 4-fraction method displayed significantly higher variability than other methods, attributed to one outlier sample. All tested ECM extraction protocols were repeated in triplicate with new starting material and the results were consistent based on performance comparisons against the 1-fraction A method (Supporting Information Figure 1). Therefore, the observed variability is likely attributed to the method rather than an error in sample preparation. The 1-fraction B method resulted in significantly fewer cellular PSMs than any other decellularization method (p=0.0052). With increased decellularization fractions, there is roughly a linear increase in total cellular PSMs (Figure 1B). In comparison of single-shot methods, in-gel and SPEED digestion result in the most cellular PSMs (p=0.0043 and 0.0027 respectively), although all single-shot extractions generate fewer cellular PSMs than the 1-fraction, 2-fraction, and 4-fraction decellularization methods when normalized to equal run time (Figure 1B).

**Figure 1.**
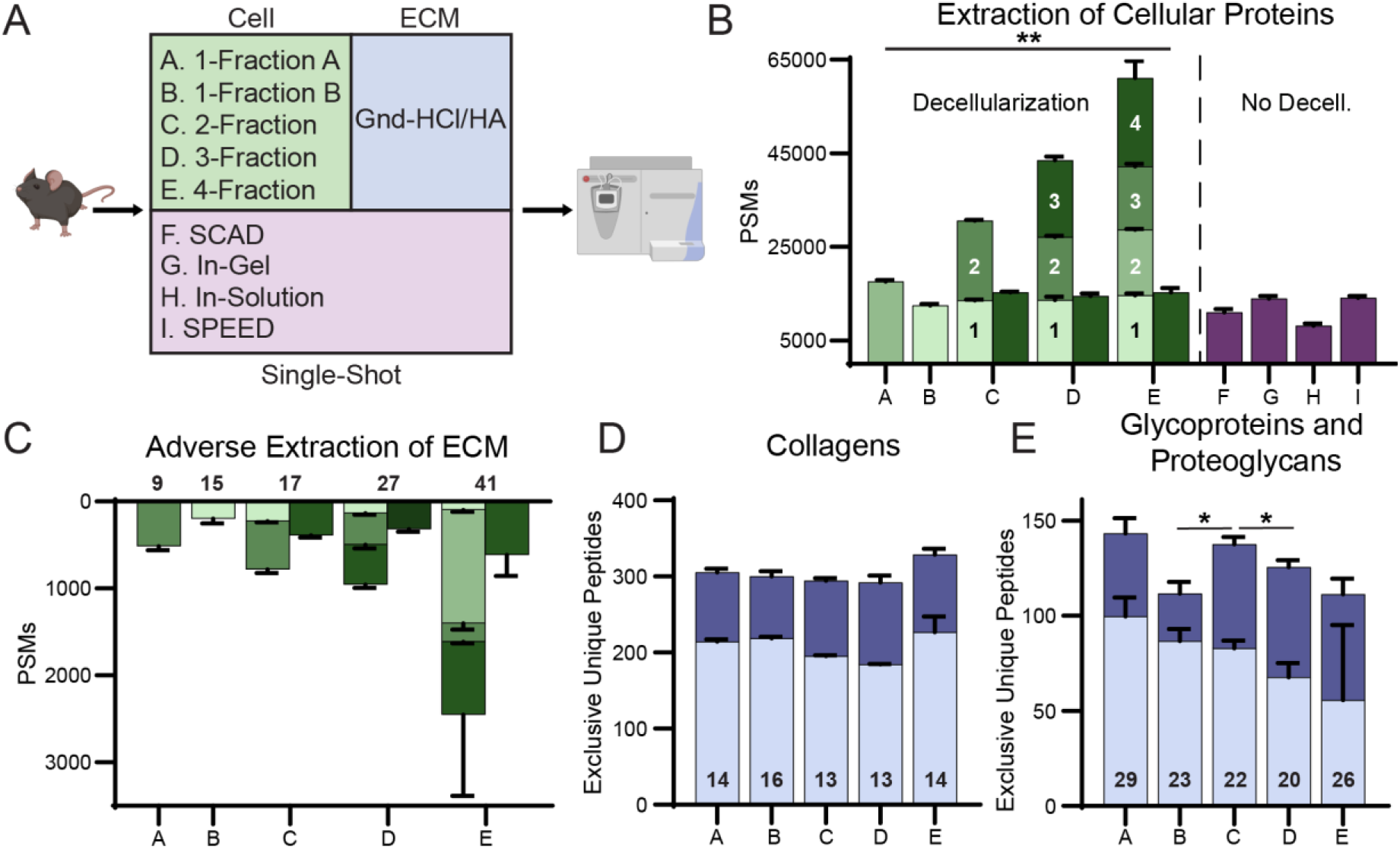
Effects of soluble protein extraction methods on extracellular matrix identification. A) Workflow of decellularization and single-shot methods used. All decellularization methods were followed by a single ECM analysis method (Gnd-HCl/HA extraction). B) Total cellular PSMs identified by each decellularization (green) and single-shot (purple) method. Multi-step methods are shown as both a sum of all fractions and the average PSMs per fraction. C) Total ECM PSMs lost during decellularization. Multi-step methods are shown as both a sum of all fractions and the average PSMs per fraction. Numbers above bars indicate the number of core matrisome proteins identified with a minimum of 2 peptides. D) Total exclusive unique collagen peptides identified in the subsequent Gnd-HCl/HA ECM extraction following each decellularization method. Light blue indicates Gnd-HCl fraction, dark blue indicates new unique peptides from HA fraction. Numbers on bars indicate the number of distinct collagen chains identified with a minimum of 2 peptides. E) Total exclusive unique peptides for proteoglycans and glycoproteins identified in the subsequent ECM extraction following each decellularization method. Numbers on bars indicate the number of ECM glycoproteins and proteoglycans identified with a minimum of 2 peptides. All bar plots present group averages with standard deviation (SD).

As stated, decellularization methods prior to ECM analysis should efficiently extract high-abundance cellular proteins while minimizing ECM extraction for subsequent analysis. Therefore, any ECM proteins removed by decellularization would be considered adverse for accuracy of subsequent ECM protein analysis. The 1-fraction B method results in the lowest number of core matrisome PSMs identified in the cellular fraction (p=0.026) (Figure 1C). This finding, in addition to the low number of cellular PSMs identified using the 1-fraction B method, indicates this method provides the mildest decellularization conditions of all the tested methods. In contrast, the 1-fraction A and 4-fraction methods extract the largest amount of core matrisome PSMs per fraction in their cellular fractions, although increased variability in the 4-fraction method prevents significance from being reached (Figure 1C). Significantly more core matrisome PSMs were identified in the second fraction of the 4-fraction decellularization method than in any individual cellular fraction from another method (136% greater, p=8.5×10^−5^), confirming that this method extracts the largest amount of ECM proteins during decellularization. For all decellularization methods, greater than 75% of PSMs of proteins classified as matrisome-associated^28^ were identified in the cellular fractions (Supporting Information Figure 2). The solubility of matrisome-associated proteins in all tested decellularization buffers supports the decision to focus on the core matrisome when evaluating ECM extraction and analysis.

**Figure 2.**
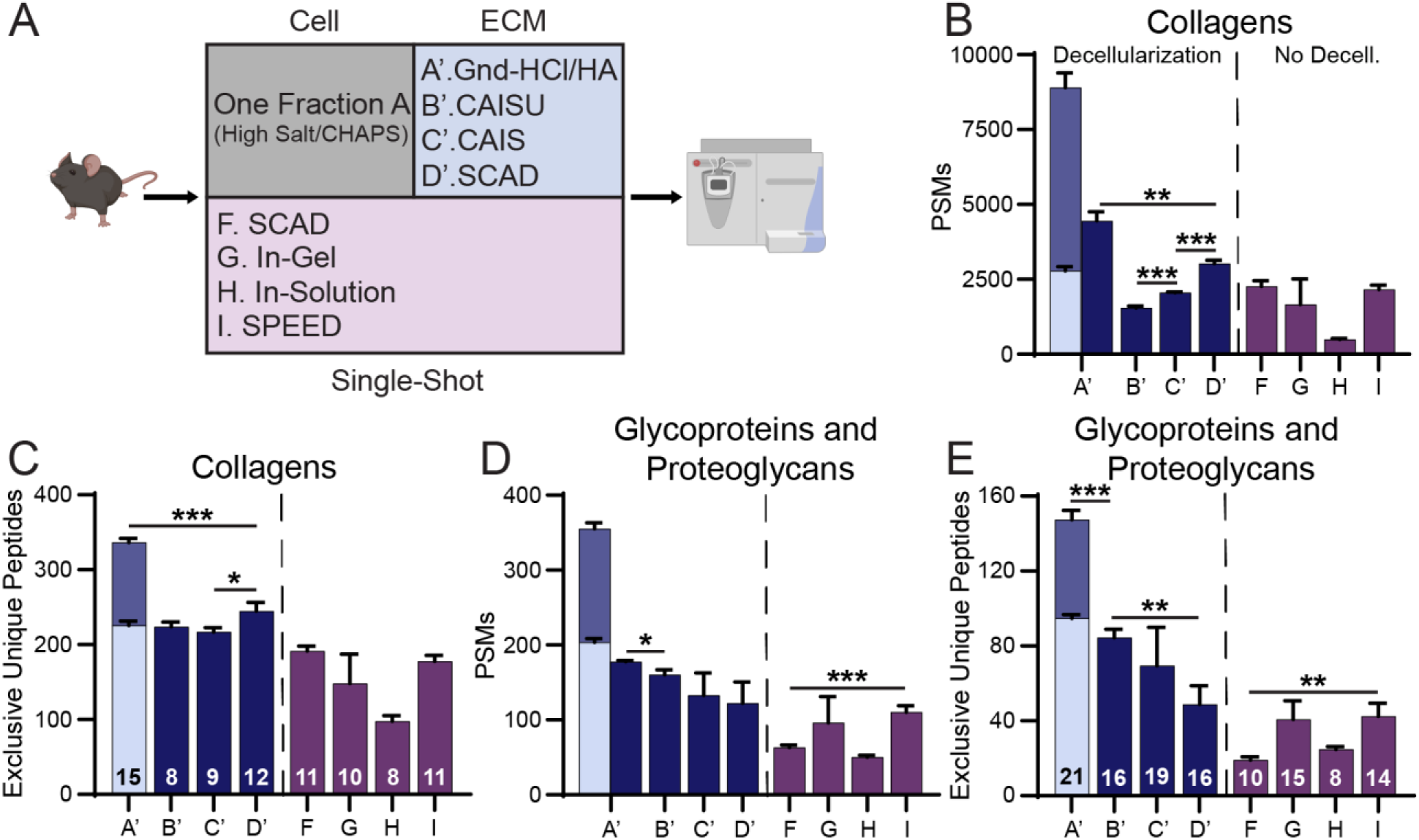
Effects of differing extracellular matrix extraction methods on uniform decellularized protein pellet. A) Workflow of ECM extraction (blue) and single-shot (purple) methods used. All ECM methods were preceded by the same decellularization. B) Total collagen PSMs identified by each method. Multi-step extraction (1’) is shown as both a sum of the two fractions and the average PSMs per fraction. Plotted as average values with SD. For Gnd-HCl/HA method, light blue indicates Gnd-HCl fraction, darker blue indicates HA fraction. C) Exclusive unique collagen peptides identified by each method. Numbers on bars represent the number of distinct collagen chains identified with a minimum of 2 peptides. D) Bar graph of glycoprotein and proteoglycan PSMs by each method. Multi-step extraction (1’) shown as both a sum of two fractions and the average PSMs per fraction. E) Exclusive unique peptides identified for glycoproteins and proteoglycans by each method. Numbers on bars represent the number of glycoproteins and proteoglycans identified with a minimum of 2 peptides.

### Evaluation of Resulting ECM Fractions Following Various Decellularization Approaches

To facilitate comparison of the remaining ECM pellets from each decellularization method, the protein composition was assessed via a single approach (the guanidine hydrochloride extraction followed by hydroxylamine digestion (Gnd-HCl/HA) method) for ECM analysis (Figure 1A, Table 1). The most effective decellularization method should result in high sequence coverage for core ECM components such as collagen, proteoglycans, and glycoproteins within the ECM fractions. The 4-fraction decellularization method resulted in 45% higher collagen PSMs within the subsequent ECM fractions than other methods (p=0.0029) (Supporting Information Figure 3A). However, it did not produce significantly more unique collagen peptides than other methods (Figure 1D). No significant difference in unique collagen peptides within the ECM fractions was identified between any tested methods (Figure 1D). The 1-fraction A (14% greater, p=0.011) and 2-fraction (9% greater, p=0.022) methods provide more unique peptides for proteoglycans and ECM glycoproteins, while the 1-fraction B and 4-fraction methods generate 12% fewer unique peptides than other methods (Figure 1E). No significant differences in ECM glycoprotein and proteoglycan PSMs were identified between any tested methods (Supporting Information Figure 3B).

**Figure 3.**
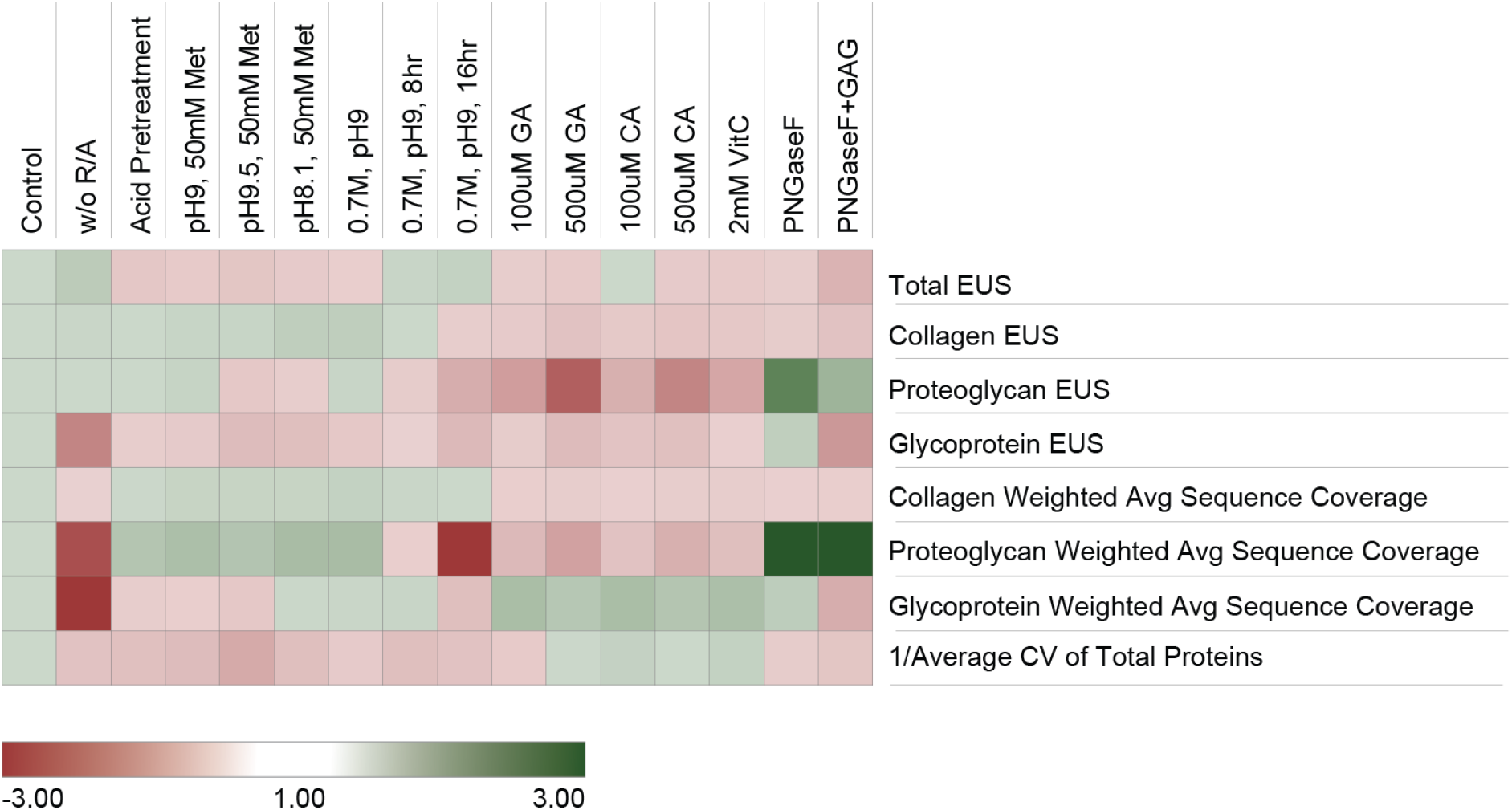
Optimization of Hydroxylamine (HA) digestion conditions. Hydroxylamine digest optimization was tested over several rounds of sample processing. Results were aggregated by comparing each digestion method to the reference sample run in parallel with treatment conditions (all n=3). Results are presented and colored as fold change in relation to reference. Reference HA digest was performed in 1M HA-HCl pH 9.0 at 37°C for 4hrs with reduction and alkylation prior to digestion. A detailed reference protocol can be found in the methods section. In the final column, GAG refers to the addition of GAG-digesting enzymes. Samples with fold change <1 are shown as negative reciprocal values.

### Evaluation of ECM Extraction and Digestion Methods

To assess ECM extraction methods on uniform starting material, a single decellularization method (1-fraction A) was performed on all samples prior to the four ECM extraction methods under evaluation (Figure 2A). Comparing all methods, the Gnd-HCl/HA method was found to result in the greatest number of identified collagen proteins, unique peptides, and PSMs even when normalized to equal run time, providing 37% more unique collagen peptides than the next best method: SCAD post-decellularization (p=0.0002) (Figure 2B,C). Additionally, Gnd-HCl/HA (p=0.0008) and SCAD post-decellularization (p=0.045) extractions generate significantly more sequence coverage of collagen proteins than the CAIS extraction method (Supporting Information Figure 4A). The CAISU and CAIS extractions resulted in similar numbers of unique collagen peptide identifications (Figure 2C). However, the CAISU method produced the lowest number of spectral matches for collagen peptides of the tested post-decellularization ECM extractions (p=0.0002) and also produced fewer collagen PSMs than the SCAD (p=0.0032) and SPEED (p=0.0029) single-shot methods despite their lack of cellular protein removal (Figure 2B). Single-shot protocols SCAD and SPEED performed similarly in all assessed metrics for collagen extraction (Figure 2B,C). Decellularization prior to the SCAD protocol offered significant improvement in identification of collagen spectral matches (p=0.0039) and unique collagen peptides (p=0.0024) (Figure 2B,C). Single-shot in-solution digestion resulted in the lowest number of collagen PSMs and unique peptides (Figure 2B). When comparing individual fractions, the HA fraction from the Gnd-HCl/HA extraction yielded the most collagen PSMs (p=0.0042) (Figure 2B).

**Figure 4.**
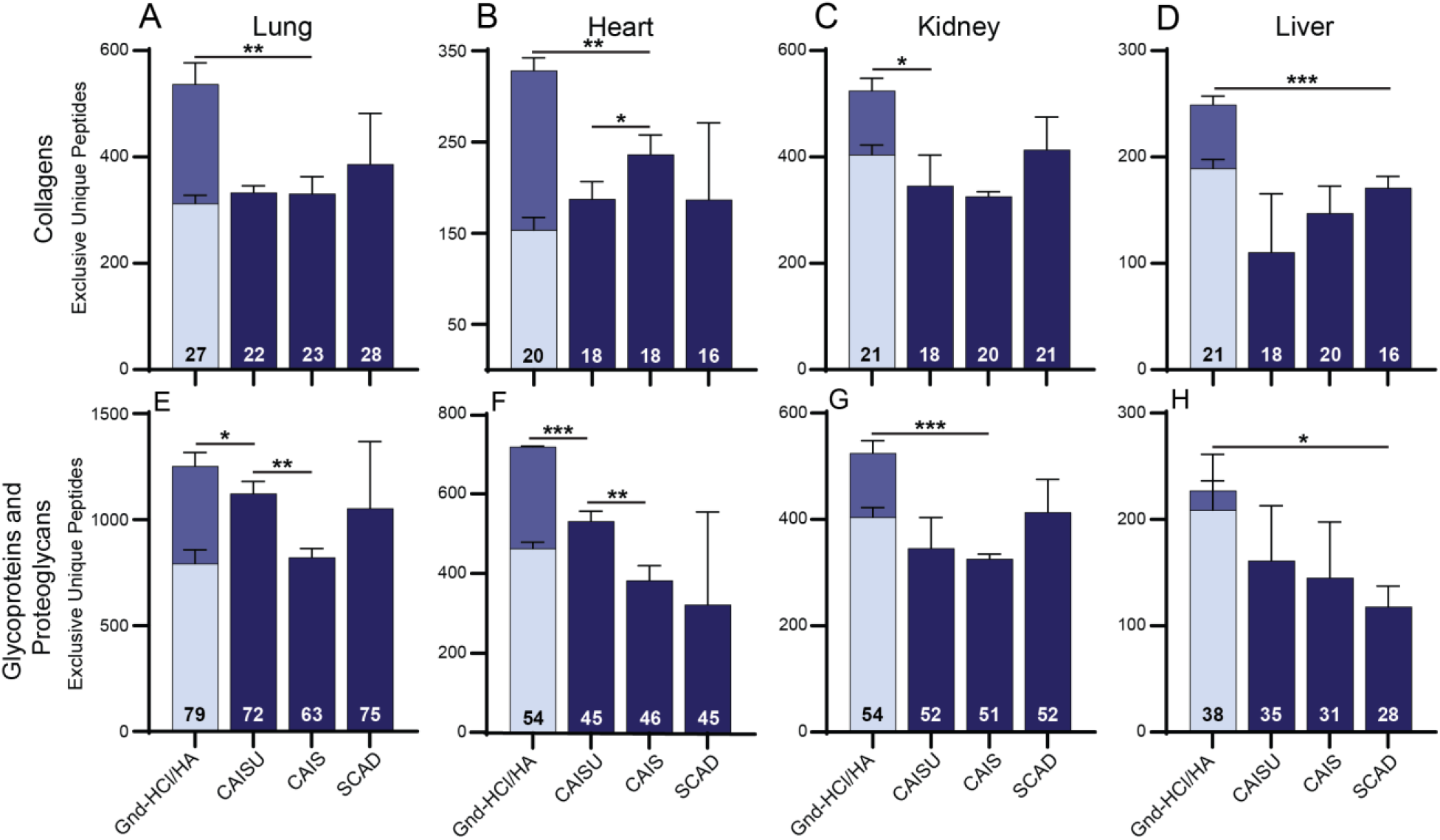
Effects of ECM extraction methods on uniformly decellularized mouse lung, heart, kidney, and liver. A-D) Total exclusive unique collagen peptides for each ECM extraction method. Numbers on bars represent the number of distinct collagen chains identified. For Gnd-HCl/HA method, light blue indicates Gnd-HCl fraction, dark blue indicates new unique peptides from HA fraction. E-H) Total glycoprotein and proteoglycan exclusive unique peptides identified by each ECM extraction method. Numbers on bars represent the number of glycoproteins and proteoglycans identified. Bars represent the average of three replicates with error bars showing standard deviation (SD).

Identification of glycoproteins and proteoglycans was also assessed to determine the best method for comprehensive core ECM analysis. The two-fraction Gnd-HCl/HA method resulted in the identification of more glycoproteins and proteoglycans (21) than any other method with 74% more unique peptides than the next best method, CAISU (p=0.0001) (Figure 2E). Additionally, the Gnd-HCl/HA extraction generated a 25% increase in glycoprotein and proteoglycan sequence coverage (p=0.024) (Supporting Information Figure 4B) and 10% more PSMs (p=0.011) than the next best method, even when normalized to equal run time (Figure 2D). SCAD extraction post-decellularization, on the other hand, produced significantly fewer unique peptides for glycoproteins and proteoglycans than both the Gnd-HCl/HA (p=0.00015) and CAISU (p=0.0051) methods (Figure 2E). SPEED extraction produced 14.5% higher glycoprotein and proteoglycan PSMs than other single-shot methods (Figure 2D), although this method identified significantly fewer unique peptides in these categories than either the Gnd-HCl/HA (p=0.00005) or CAISU (p=0.00096) extractions (Figure 2E) and fewer glycoproteins and proteoglycans than any post-decellularization method (Figure 2E). When comparing individual fractions, the first fraction generated by the Gnd-HCl/HA method generates the greatest number of unique glycoprotein and proteoglycan peptides (p=0.023) (Figure 2E). Overall, utilizing a two-fraction Gnd-HCl/HA extraction provides significantly greater identification of collagen, proteoglycan, and glycoprotein peptides than all other tested methods.

ECM extraction methods which utilize urea have the potential to induce carbamylation at lysine and arginine residues, as well as peptide N-termini. This modification can inhibit trypsin digestion and increase sample heterogeneity, convoluting data analysis and subsequent quantification.^34^ In our analysis, we identify averages of 763, 641, and 1523 carbamylated peptides per run in the CAISU, CAIS, and SCAD methods, respectively, compared to an average of 34 carbamylated peptides identified across the two fractions of the Gnd-HCl/HA method (Supporting Information Figure 5A). While the SCAD method induces more carbamylation than the other enzymatic extractions, all methods that utilize urea in an extraction buffer induce this modification extensively, likely contributing to deficits in ECM identification. On the other hand, hydroxylamine digest induces oxidation over other methods, where an average of 5954 oxidized peptides per run were identified in the HA fraction of the Gnd-HCl/HA method, compared to averages of 1702, 2290, and 3286 oxidized peptides in the CAISU, CAIS, and SCAD extracts, respectively (Supporting Information Figure 5B). While oxidation does not inhibit peptide cleavage, it can convolute analysis and protein quantification.

### ECM Extraction and Digestion Method Optimization

The 2-fraction Gnd-HCl/HA extraction method was used for further optimization of ECM coverage, based on the above findings. Oxidative modifications observed with the use of hydroxylamine digestion^4^ can convolute data analysis and have the potential to reduce both the quantity and quality of protein identifications. On the other hand, if oxidation of Met residues is pushed toward completion without inducting other significant oxidations, this would have an advantageous effect on these metrics. In order to optimize this step, iterative changes to the hydroxylamine digestion protocol were explored with the goal of reducing unwanted modifications and ultimately increasing ECM protein identification. A reference protocol, as used in the previous section, was analyzed in parallel with each group of optimization conditions. Briefly, the reference protocol involves digestion with 1M HA, pH 9.0 for 4 hours at 37°C with reduction and alkylation (R/A) prior to digestion. The previously published Gnd-HCl/HA method^4^ did not include R/A due to the lack of reducible disulfide bonds in collagen. However, R/A provided a 60% improvement in unique glycoprotein peptide identifications (p=0.0088), including a 10-fold increase in PSMs for fibrillin-1 (Supporting Information Figure 6) and improved overall CVs (Figure 3). Therefore, R/A prior to HA digest was used as the reference method for the subsequent method optimization experiments. The second variable tested was pre-treatment of the pellet with 0.2% FA, as suggested by Bornstein and Bailan,^35^ which showed no significant benefit and increased the overall CV of ECM measurements (Figure 3).

We also evaluated the effects of changes in pH, HA concentration, digestion time, and addition of an antioxidant (50mM methionine) on identification of ECM components. Most tested variations led to slight improvements in collagen PSMs but higher CVs and fewer total PSMs (Figure 3). Of note, pH 8.1 with 50mM methionine improved the number of identified collagen PSMs. However, decreases in total and glycoprotein PSMs, as well as increased CV, make the method less suitable than the reference method unless maximizing total collagen PSMs is an objective of the analysis. In addition, 0.7M HA digestion for 16 hours performed significantly worse for all scoring metrics (Figure 3). This was consistent with previous findings where extended digestion times result in extensive peptide modification.^4^

50mM methionine showed potential improvements in collagen PSMs and coverage over conditions without antioxidant. Therefore, the effects of more potent antioxidants including caffeic acid (CA), gallic acid (GA), and ascorbic acid (VitC) at varying concentrations on HA digestion were tested. All antioxidants showed significant improvement in sequence coverage of glycoproteins but not of proteoglycans or collagens (Figure 3). This is coupled with a significant drop in proteoglycan PSMs in comparison to the reference method. A final variable explored was deglycosylation, as improved identification of ECM glycoproteins and proteoglycans has been reported using PNGase F to remove N-linked glycans,^20,36^ as well as GAG-digesting enzymes chondroitinase ABC and heparinase II.^36–38^ The addition of PNGaseF slightly improved proteoglycan PSMs and sequence coverage over the reference method but also generated slight decreases in collagen PSMs and sequence coverage. PNGase F digestion significantly improved glycoprotein coverage (p=0.017) but did not improve glycoprotein PSMs (Figure 3). The addition of GAG removal enzymes (chondroitinase ABC and heparinase II) offered a significant improvement in proteoglycan coverage (p=0.012) but not PSMs. Improvement in proteoglycan identification was coupled with decreases in collagen (p=0.0044) and glycoprotein (p=0.0002) PSMs, causing GAG removal to result in overall worse ECM characterization than the reference method (Figure 3). When comparing deglycosylation (PNGase F) of both the Gnd-HCl and HA fractions, in general similar trends were observed when compared to the reference method for PSMs and sequence coverage of proteoglycans (Supporting Information Figure 7). However, improvement of proteoglycan coverage with the addition of PNGase F was highly significant (p=0.00082) when the Gnd-HCl fraction was taken into account.

### Evaluation of ECM Extraction and Digestion Methods on Isolated Organs

All previous comparisons of ECM extraction methods were performed on an early generation Orbitrap mass spectrometer (MS) due to the widespread use of this analytical platform and accessibility in core facilities. In order to perform more in-depth comparisons of ECM extraction methods and to determine if conclusions drawn from WMP hold true for organ specific analysis, extractions of lung, heart, kidney, and liver were analyzed on a modern, trapped ion mobility-time of flight MS system (Figure 4). Organs were chosen based on their widespread study in biomedical research and their varying ECM compositions. Organ samples were prepared using the 1-fraction A decellularization followed by the four tested ECM extraction methods.

Consistent with our findings from WMP analysis, the Gnd-HCl/HA method provides more exclusive unique peptides for collagens in the lung (39% greater), heart (38% greater), kidney (26% greater), and liver (46% greater) than any other method (Figure 4A-D). Additionally, the Gnd-HCl/HA method performs similarly to, or better than, other methods in terms of glycoprotein and proteoglycan identification. The Gnd-HCl/HA method provided significantly more exclusive unique peptides than the next best method in lung (11% greater, p=0.026) and heart (32% greater, p=0.0004) samples, but statistical significance was not reached in other organ comparisons (Figure 4E-H). Also consistent with results from WMP, the CAISU method performs better for identification of glycoproteins and proteoglycans across all organs than either the CAIS or SCAD extraction methods (Figure 4E-H).

## Discussion

Based on the significant role that the structural matrix plays in shaping cellular phenotype^39,40^ and, likewise, the influence of cell phenotype on stromal and matrix composition,^41^ optimized proteomic methods for ECM characterization are needed. In general, accurate tissue characterization requires analysis of the extracellular matrix, yet this class of proteins is under-represented in tissue proteomic datasets. Several proteomic methods have been developed to improve characterization of the ECM over the last two decades. Key elements of these protocols involve 1) removal of cellular material to create an ECM enriched fraction, primarily using differential detergent extraction and 2) solubilization and efficient digestion of the resulting ECM. Various published decellularization methods were assessed for their ability to extract cellular proteins while leaving ECM proteins behind for analysis in subsequent fractions. The 1-fraction A and 4-fraction methods were found to remove more cellular proteins from the starting material than other tested methods. If cellular characterization and solubility profiling of cellular components are desired alongside ECM characterization, multi-fraction methods provide more cellular protein coverage when all fractions are analyzed. Also, of no surprise, single-shot methods result in fewer cellular PSMs than nearly all decellularization methods, with lower identifications due to the increased complexity and dynamic range of the extracted proteins, in part due to partial ECM extraction.

The largest amount of ECM proteins was extracted during decellularization using the 4-fraction method, evidenced by the high number of core matrisome protein and PSM identifications in the cellular fractions. Extraction of ECM proteins in the cellular fraction is undesirable due to difficulty detecting ECM proteins of interest against the background of abundant cellular protein. Additionally, when the 4-fraction method is used for decellularization prior to ECM extraction, the cellular fractions are often not analyzed via MS,^20,42^ causing all ECM proteins present in these fractions to be lost and ultimately resulting in quantitative error and potentially fewer identified core matrisome proteins within the ECM fractions. In our experience, to analyze these cellular fractions via LC-MS/MS additional detergent removal steps must be performed to avoid instrument contamination. Although the 1-fraction B method removes the least amount of ECM during decellularization, it does not efficiently extract cellular proteins, thus limiting subsequent ECM characterization.

After decellularization, the resulting pellets were processed using a single method for direct comparison of the resulting ECM fractions. Similar collagen PSMs were identified in the 1-fraction A, 2-fraction, and 3-fraction methods, while the 4-fraction method resulted in significantly more collagen PSMs but fewer collagens and similar collagen sequence coverage to other methods. This is due to the high stringency of the 4-fraction decellularization method, extracting most non-collagen proteins in earlier fractions and allowing more sequencing time to be devoted to a subset of collagen peptides in the final ECM fraction. If optimal collagen signal is desired, performing 4-fraction decellularization before ECM extraction can provide greater collagen spectral matches but not necessarily higher sequence coverage of collagen proteins. Additionally, the 4-fraction method generates significantly less sequence coverage of ECM glycoproteins and proteoglycans, hampering its overall characterization of core matrisome proteins. The 2-fraction decellularization method, on the other hand, provided the greatest coverage of glycoproteins and proteoglycans within the ECM fractions, making this the method of choice when these protein classes are a priority.

Total PSMs, ECM PSMs, and method precision have been discussed above. These evaluation metrics along with time, ease, and MS compatibility are shown in Table 2. Each category was ranked on a scale of 1-10 with 1 being the lowest possible score and 10 being optimal (ranking criteria are further defined in methods). MS compatibility was ranked based on the use of MS-incompatible detergents throughout the course of the method. Acetone precipitation or a detergent removal column can be utilized within any protocol to reduce risk of MS contamination, but these steps can generate additional cost, variability, and protein loss. Methods that utilize SDS and NP-40, such as the 3-fraction method, can cause residual contamination during MS analysis of subsequent ECM fractions and require additional pellet washing or acetone precipitation of the ECM fractions. The ease criterion is a ranking of the relative ease of each method in comparison to the other protocols, considering the number of processing steps and potential for increased analytical CV. The 4-fraction method displayed higher variability than other methods across two triplicate rounds, likely due to the large number of buffer exchanges required during processing (10) compared to other methods. As this table shows, there is not one method that is superior in all categories. Instead, a method should be selected specifically based on project aims, constraints, and proteins of interest.

**Table 2.** Comparison of decellularization methods for ECM enrichment. Decellularization (green) and single-shot (purple) methods are ranked in each category from one to ten - one being totally insufficient and ten being optimal. Total score is calculated as a weighted average of all assessed categories (described in methods).

| Method | Time | Ease | MS Compatability | Total Protein IDs | Precision | ECM in Cell Fraction | Total Score |
| --- | --- | --- | --- | --- | --- | --- | --- |
| 1-Fraction A | 8 | 9 | 7 | 8 | 9 | 8 | 82 |
| 1-Fraction B | 3 | 9 | 6 | 7 | 8 | 9 | 74 |
| 2-Fraction | 9 | 8 | 6 | 8 | 9 | 7 | 79 |
| 3-Fraction | 8 | 7 | 6 | 9 | 8 | 7 | 77 |
| 4-Fraction | 6 | 6 | 6 | 9 | 6 | 6 | 68 |
| SCAD | 6 | 8 | 8 | 6 | 8 | N/A | 71 |
| In-Gel | 7 | 7 | 8 | 7 | 6 | N/A | 69 |
| In-Solution | 9 | 9 | 9 | 5 | 9 | N/A | 78 |
| SPEED | 9 | 7 | 9 | 7 | 9 | N/A | 81 |

The second key element of an effective ECM proteomic approach is efficient digestion of matrix proteins. This task is hampered by the relative protease resistance of the ECM. Four published ECM extraction methods were evaluated on both WMP and isolated organ samples following uniform decellularization. While it is impractical to test all published methods and extensively test all variables, these four methods cover the major ECM-based methods that are currently used. As stated above, the two-step Gnd-HCl/HA method identified the greatest number of proteins, PSMs, and unique peptides from all core matrisome categories in analysis of both WMP and isolated organs. While the Gnd-HCl fraction of the Gnd-HCl/HA extraction alone performs similarly to other tested methods, the following HA digest provides 355 additional unique core matrisome peptides on average across analyzed organs, leading the 2-fraction HA digest method to provide improved core matrisome characterization over other tested methods. This, combined with the higher sequence coverage of collagens, glycoproteins, and proteoglycans, makes the Gnd-HCl/HA extraction the recommended method for obtaining more in-depth coverage of the ECM, despite the additional time it requires. The proportion of ECM peptides which are uniquely identified by the HA digest in the Gnd-HCl/HA method varies greatly between tested organs, with 40% of ECM peptides uniquely identified in the HA fraction of heart samples but only 15% in the HA fraction of kidney. These differences reflect varying relative ECM protein abundance, association, and crosslinking - attributes which are often altered during development and disease progression.

Of the enzymatic ECM extractions, SCAD after decellularization provides a greater number of identified collagens and more unique collagen peptides than the CAIS or CAISU methods in WMP analysis. The CAISU method, on the other hand, yielded more unique glycoprotein and proteoglycan peptide identifications and greater sequence coverage than other enzymatic methods in all samples. Therefore, enzymatic methods should be chosen depending on the ECM protein categories of interest when only one ECM fraction is analyzed.

When comparing faster, single-shot methods, SCAD and SPEED generated significantly higher PSMs for collagens than the CAISU method, even though these methods lack decellularization steps to remove cellular contaminants. SPEED also provides the greatest number of unique peptides from glycoproteins and proteoglycans of the tested single-shot methods. SCAD performs poorly in this matrisome category, revealing that SPEED is the best performing single-shot method for overall ECM coverage. SPEED is a favorable alternative to longer ECM extractions that provides good ECM coverage and unique peptide IDs when higher throughput ECM analysis is desired, and multiplexing is not available or otherwise used.

A condensed evaluation of all ECM extraction and single-shot methods can be found in Table 3. Some of the criteria for evaluation have already been discussed above. This table also addresses the time needed to perform each extraction and the relative ease of each protocol. There is no one perfect choice for an ECM extraction method but these evaluation metrics should help guide researchers to a method of choice for a given set of objectives.

**Table 3.**
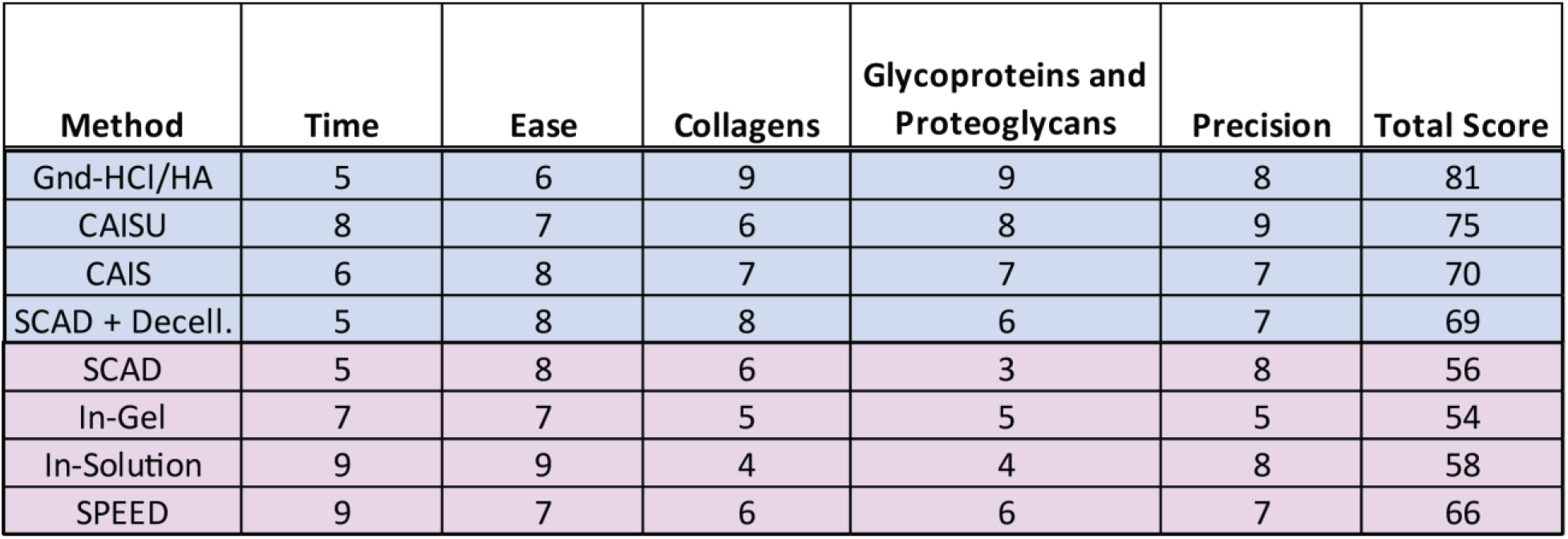
Comparison of ECM extraction/digestion methods. ECM extraction (blue) and single-shot (purple) methods are ranked in each category from one to ten - with one being totally insufficient and ten being optimal. Total score is calculated as a weighted average of all assessed categories (described in methods).

| Method | Time | Ease | Collagens | Glycoproteins and<br>Proteoglycans | Precision | Total Score |
| --- | --- | --- | --- | --- | --- | --- |
| Gnd-HCl/HA | 5 | 6 | 9 | 9 | 8 | 81 |
| CAISU | 8 | 7 | 6 | 8 | 9 | 75 |
| CAIS | 6 | 8 | 7 | 7 | 7 | 70 |
| SCAD + Decell. | 5 | 8 | 8 | 6 | 7 | 69 |
| SCAD | 5 | 8 | 6 | 3 | 8 | 56 |
| In-Gel | 7 | 7 | 5 | 5 | 5 | 54 |
| In-Solution | 9 | 9 | 4 | 4 | 8 | 58 |
| SPEED | 9 | 7 | 6 | 6 | 7 | 66 |

Based on the favorable results obtained using the two-step Gnd-HCl/HA method, we performed comparisons aimed to improve upon this method. Reduction and alkylation prior to hydroxylamine digestion was the first change tested. Based on superior initial results, HA digestion with prior reduction and alkylation became the reference method for subsequent optimization. Most tested variables resulted in both improved and reduced individual ECM category performance. Given the fact that most alterations added an additional cost or step, which increases potential variability, for most projects the published HA digestion method^4^ with added reduction and alkylation prior to HA digestion is recommended. The one exception is the addition of PNGaseF for proteoglycan coverage. Somewhat surprisingly, there is a clear increase in proteoglycan PSMs upon addition of PNGaseF but no improvement in identification of collagens or ECM glycoproteins. It is important to note that the use of PNGaseF adds extra cost to each sample preparation. In addition, the improvement in proteoglycan coverage but not PSMs suggests that some extracted glycopeptides remained unidentified by database searching until deglycosylated by PNGaseF. The addition of PNGaseF could be a worthwhile investment if optimizing proteoglycan coverage is a priority.

The first set of analysis was performed using an older generation orbitrap instrument that is likely to be encountered by many groups interested in the analysis of ECM through collaboration or core facilities. While this analytical platform offers high mass accuracy and resolution which are valuable for reducing false positives, the sensitivity and scan speed does not match more modern instrumentation. Additionally, WMP was used for initial testing due to the high complexity of the sample, allowing for comparison of methods without biasing results toward a specific application. However, proteomics studies are generally performed using isolated tissue samples, their substructures, or micro-dissected regions. As a result, we also evaluated the chosen ECM extraction methods on isolated organs using a modern timsTOF mass spectrometer to assess method performance on real-world samples using a state-of-the-art instrument. In general, comparisons between methods revealed during WMP testing were consistent with those observed in analysis of individual organs. While method comparisons are mostly consistent across the tested organs, further method optimization may be required to obtain high-quality ECM characterization for organs with very unique compositional profiles (e.g. bone). Here, we evaluated extraction and digestion of tissues in a non-limiting sample regime (1-5 mg dry weight). However, we have also had success applying our ECM extraction method to human tissue samples (<10μg) from laser-capture microdissection (LCM) extraction (Supporting Information Figure 8) with good ECM coverage, extending these techniques for spatial analysis of the ECM.

Of the over 200 proteins defined as “core matrisome” within the MatrisomeDB,^28^ 138 were identified across the 4 analyzed organs. Within the ECM, collagen is both highly abundant and highly modified. Increased fractionation of ECM samples may reveal differences in identification of low-abundance ECM proteins between methods which were not addressed in this study. All data was acquired using data-dependent acquisition because it is the most accessible acquisition method and does not require generation of a spectral library to achieve high-quality results. However, this allows for stochastic sampling of low abundance precursor species and generates more missing values. This limitation could be diminished by using data-independent acquisition, as has been previously demonstrated for ECM analysis.^43^ However, the high complexity of post-translational modifications commonly encountered on ECM peptides (more than 5 modifications per peptide with various positional isomers) presents a significant, largely unresolved issue with search routines. This also presents unique opportunities for the use of ion mobility for further resolution and characterization of these complex peptide species.

The choice of decellularization method should be based on the objectives of a given project. If minimizing sample processing time and ECM extraction are priorities, the 1 fraction A method is recommended. However, if deeper proteome coverage or solubility profiling of cellular components is desired, the 2-fraction or 3-fraction decellularization methods with analysis of all produced fractions are recommended. Deeper cellular coverage could also be provided by performing the 1 fraction A decellularization followed by offline fractionation prior to MS analysis with less potential for variability compared to these methods. For optimal ECM coverage, the Gnd-HCl/HA extraction protocol is recommended as it produces the greatest number of PSMs and highest sequence coverage of core matrisome proteins. Disease progression and aging have been shown to increase the resistance of the insoluble ECM to extraction,^44^ increasing the necessity of effectively extracting the insoluble ECM via chemical digestion. Additionally, the Gnd-HCl/HA extraction produces two separate ECM fractions which can be analyzed independently, further increasing ECM identifications and allowing for assessment of ECM solubility. Alterations in protein solubility can provide important information regarding disease progression^45,46^ which cannot be derived from single-fraction abundance measurements alone. Therefore, the Gnd-HCl/HA method is the method of choice for optimized ECM analysis. For a single-shot method, the SPEED method is the method of choice due to its ability to provide moderately high coverage of both core matrisome and cellular components using a single MS run. However, throughput can also be increased using multiplexing reagents and faster acquisition times available on modern MS instrumentation.

Mass spectrometry has increasingly played a central role in biomolecular characterization of tissues for the study of development, health, and disease. While the instrumentation, data acquisition routines, and bioinformatic tools facilitating proteomics workflows are consistently improving, less attention has been given to improving sample preparation methods and ensuring that all relevant material from a sample is analyzed. Without optimized sample preparation methods, a significant portion of the tissue proteome will be missed. ECM proteins will remain insoluble and studies will not achieve characterization of a protein fraction that is largely responsible for the underlying cell phenotype and biomechanics of a tissue. The methods evaluated and optimized here should help facilitate studies of tissue microenvironments in development, disease progression and aging.

## ASSOCIATED CONTENT.

## Supporting Information

The following files are available.

Supplementary Figures (.docx)

Supplementary Data – Exclusive Unique Peptides

(SupplementaryData_ExclusiveUniquePeptides.xlsx)

Supplementary Data – Total Peptide Spectral Matches

(SupplementaryData_TotalPeptideSpectralMatches.xlsx)

The mass spectrometry proteomics data have been deposited to the ProteomeXchange Consortium via the MassIVE partner repository with the data set identifier PXD021778. Data available upon request – contact corresponding author for access.

## AUTHOR INFORMATION

### Author Contributions

The manuscript was written through contributions of all authors. All authors have given approval to the final version of the manuscript. ‡MCM and LRS contributed equally.

## ACKNOWLEDGMENT

We would like to thank the numerous students, technicians and collaborators that have evaluated, utilized, and provided feedback on these methods throughout the years. Workflow figures throughout the publication were created with BioRender.com.

## ABBREVIATIONS

ABC: Ammonium bicarbonate
ACN: Acetonitrile
CA: Caffeic acid
CAIS: Chaotrope-assisted in-solution digest
CAISU: Chaotrope-assisted in-solution digest with ultrasonication
CHAPS: 3-[(3-cholamidopropyl)dimethylammonio]-1-propanesulfonate
CV: Coefficient of variance
DOC: Sodium deoxycholate
DTT: Dithiothreitol
ECM: Extracellular matrix
EDTA: Ethylenediaminetetraacetic acid
FA: Formic acid
FDR: False discovery rate
GA: Gallic acid
GAG: Glycosaminoglycan
Gnd-HCl: Guanidine hydrochloride
HA: Hydroxylamine hydrochloride
HEPES: 4-(2-hydroxyethyl)-1-piperazineethanesulfonic acid
IAM: Iodoacetamide
K2CO3: Potassium carbonate
KCl: Potassium chloride
LC-MS/MS: Liquid chromatography tandem mass spectrometry
MgCl_2_: Magnesium chloride
MS: Mass spectrometry
NaCl: Sodium chloride
NaOV: Sodium orthovanadate
NP-40: Nonidet-P40
PI: Protease Inhibitor
Pipes: Piperazine-N,N′-bis(2-ethanesulfonic acid)
PSM: Peptide spectral match
R/A: Reduction and alkylation
SCAD: Surfactant and chaotropic agent assisted sequential extraction/on-pellet digestion
SDS: Sodium dodecyl sulfate
SPEED: Sample preparation by easy extraction and digestion
TFA: Trifluoroacetic acid
Tris-HCl: Tris(hydroxymethyl)aminomethane hydrochloride
VitC: Ascorbic acid
WMP: Whole mouse powder

