## Supplemental Figures for "Evaluation and refinement of sample preparation methods for extracellular matrix proteome coverage"

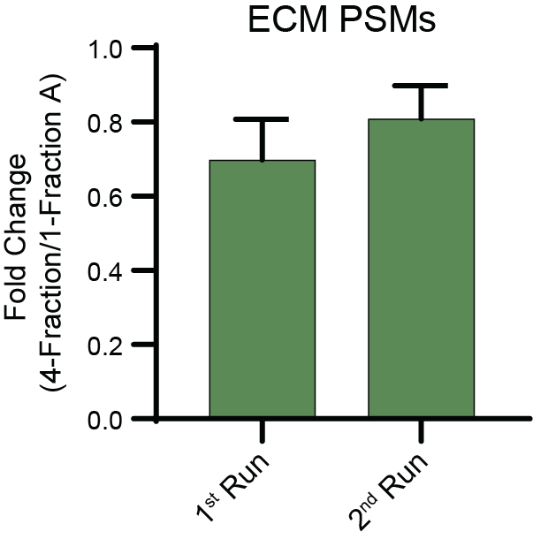

**Supporting Information Figure 1. Variability of four-fraction decellularization method across two replicate analyses.** Average fold change comparing ECM PSMs identified in subsequent ECM fractions from the four-fraction and one-fraction A methods. First run refers to triplicate analysis of samples that were run during the first round of testing. Second run refers to replicates of the four-fraction and one-fraction A methods performed using new starting material and run at a later date. Bar graph represents group averages with standard deviation (SD).

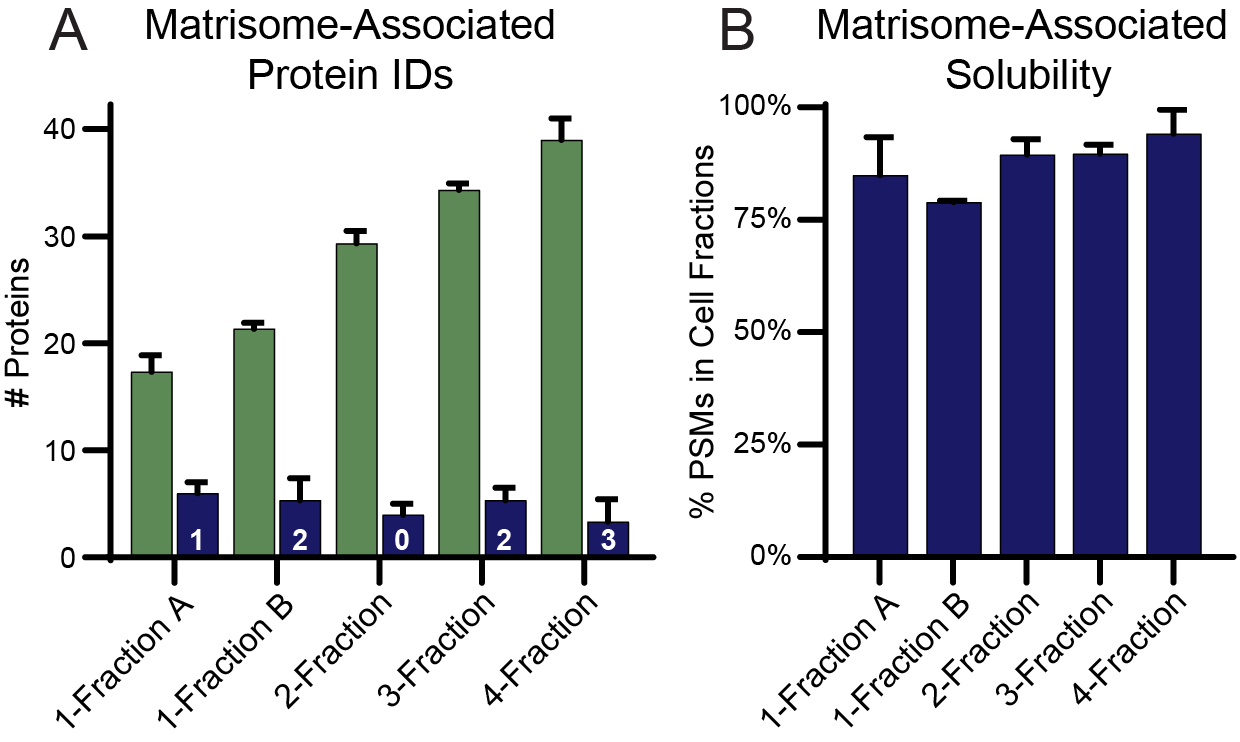

**Supporting Information Figure 2. Solubility profiling of matrisome-associated proteins.** A) Number of matrisome-associated proteins identified in the cell (green) and ECM (blue) fractions from each decellularization method. Numbers on bars indicate the number of additional matrisome-associated proteins uniquely identified in the ECM fractions. B) Average percent of total (cell + ECM) PSMs for matrisome-associated proteins identified in the cell fractions from each decellularization method. Only proteins identified in at least one fraction of every replicate are included.

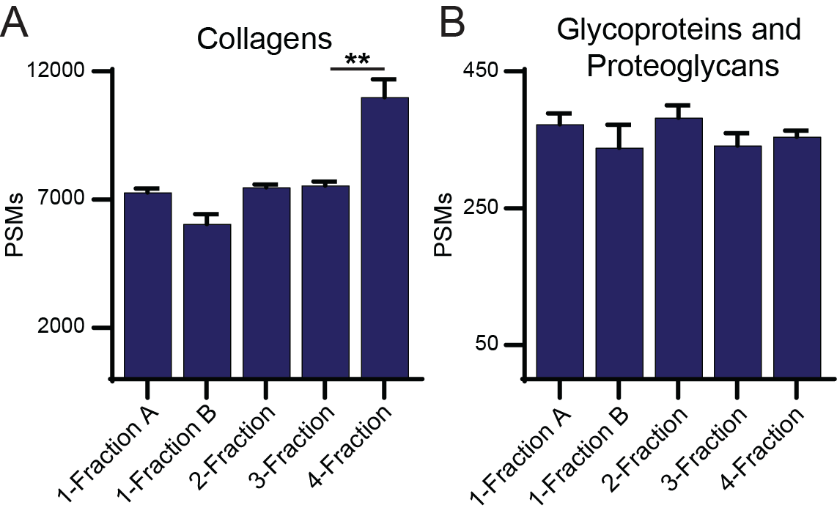

**Supporting Information Figure 3. Effects of decellularization methods on subsequent extracellular matrix (ECM) characterization.** A) Total collagen peptide spectral matches (PSMs) identified in the subsequent ECM extraction following each decellularization method. B) Total proteoglycan and glycoprotein PSMs identified in the subsequent ECM extraction following each decellularization method. All bar plots present group averages with standard deviation (SD).

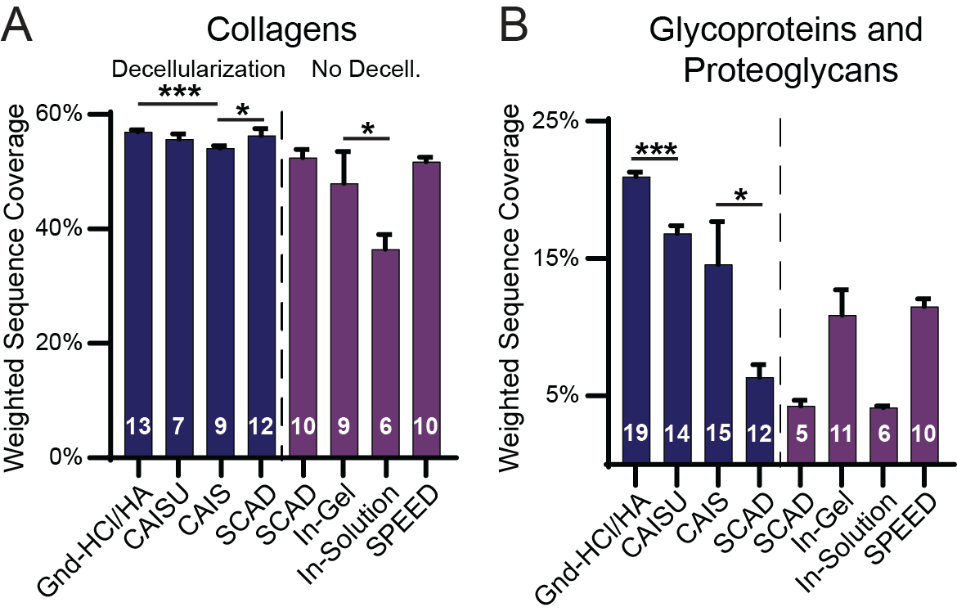

**Supporting Information Figure 4. Effects of ECM extraction on collagen and proteoglycan sequence coverage.** A) Weighted average of sequence coverage for all collagens by each method. Numbers on bars represent the number of unique collagen chains identified in at least two of three method replicates. B) Weighted average sequence coverage for all glycoproteins and proteoglycans by each method. Numbers on bars represent the number of glycoproteins and proteoglycans identified in at least two of three method replicates.

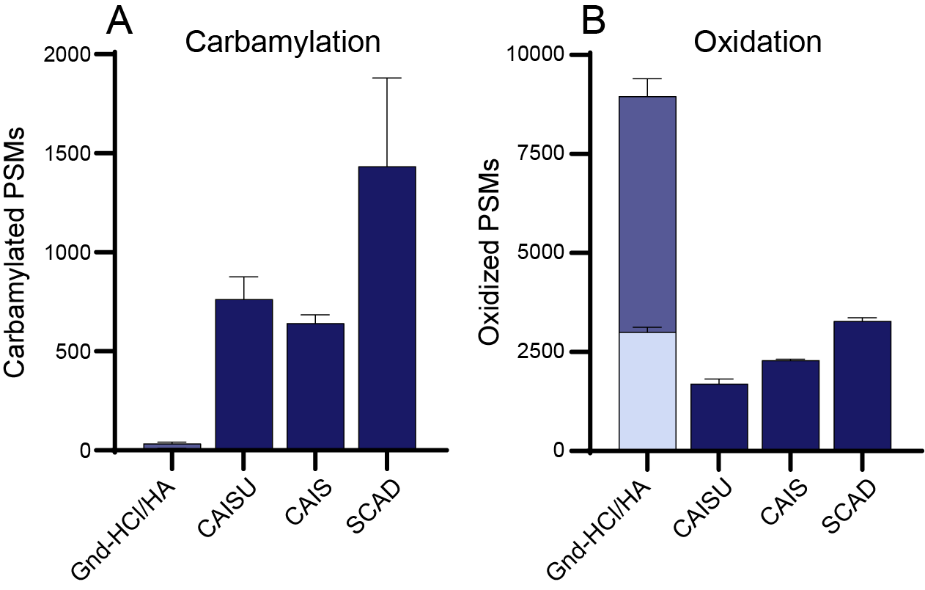

**Supporting Information Figure 5. Effects of ECM extraction method on peptide carbamylation and oxidation.** Total PSMs identified by each method with at least one carbamylation (A) or oxidation (B). Bar plot represents group averages with standard deviation (SD).

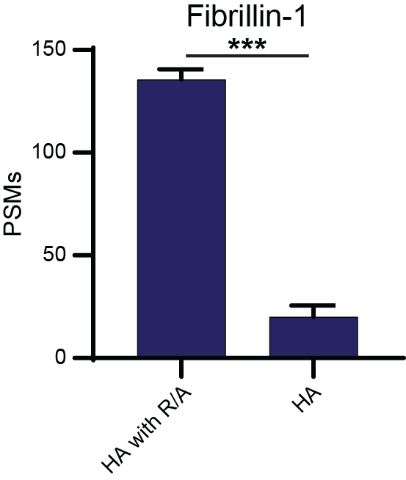

**Supporting Information Figure 6. Effects of reduction and alkylation on fibrillin-1 PSMs.** Total fibrillin-1 PSMs identified in the HA fraction with and without reduction and alkylation prior to HA digestion. Bar plot represents group averages with standard deviation (SD).

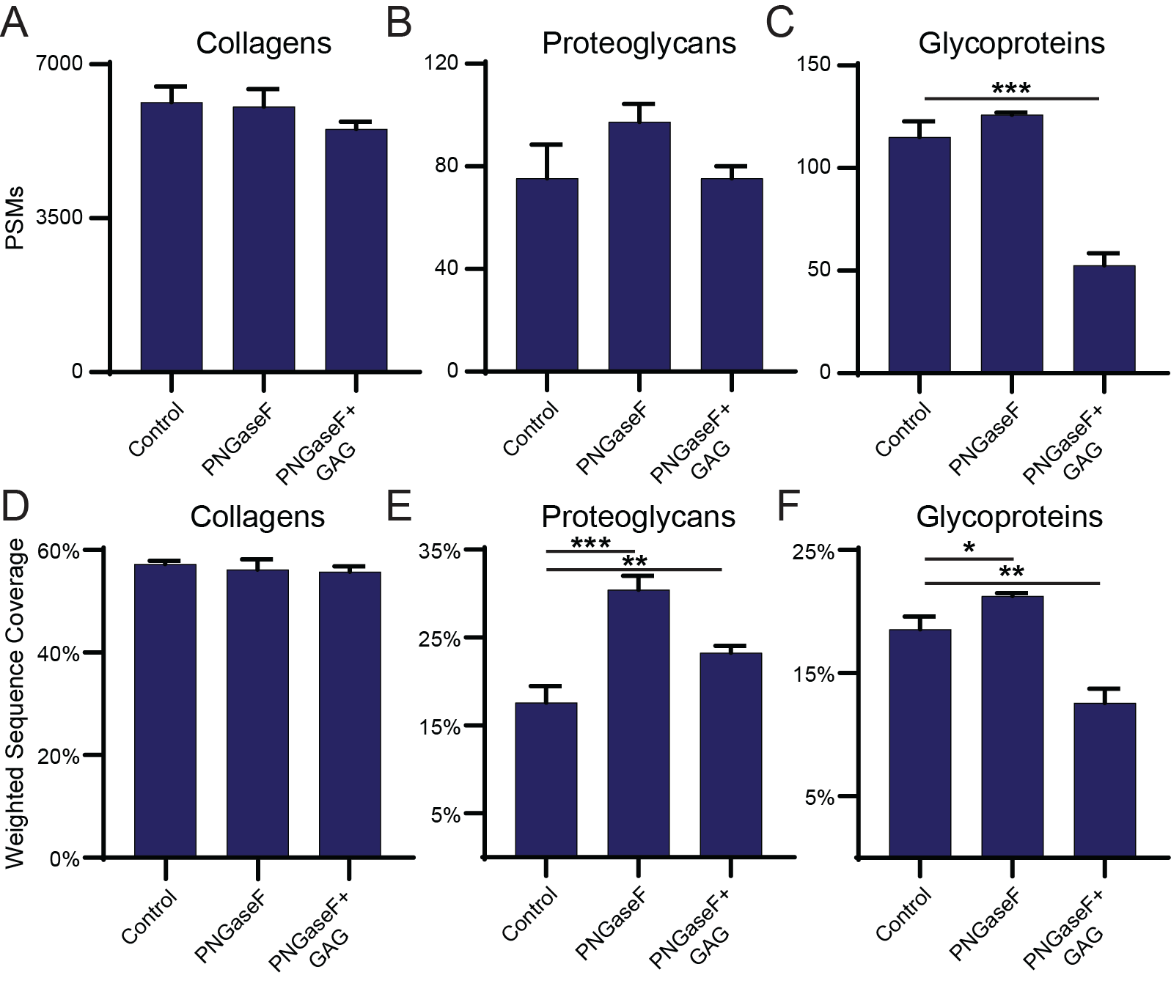

**Supporting Information Figure 7. Effect of deglycosylation on identification of ECM proteins.** A-C) Bar graphs of collagen, proteoglycan, and glycoprotein PSMs in the Gnd-HCl and HA fractions across the three methods, respectively. D-F) Bar graphs of collagen, proteoglycan, and glycoprotein weighted average sequence coverage in the Gnd-HCl and HA fractions across the three methods, respectively.

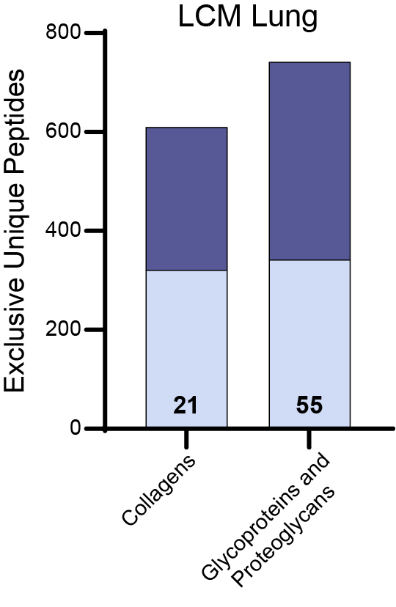

**Supporting Information Figure 8. Identification of ECM proteins in laser capture microdissection (LCM) sample from a human lung.** Total exclusive unique peptides for collagens, glycoproteins and proteoglycans identified in the Gnd-HCl (light blue) and HA (dark blue) fractions from the Gnd-HCl/HA method. Numbers on bars represent the number of identified proteins from each category. In brief, LCM lung sample was decellularized using the 1-fraction A method followed by the Gnd-HCl/HA ECM extraction method. All fractions were digested prepared for MS as described in methods.
